# An all-solid-state heterojunction oxide transistor for the rapid detection of biomolecules and SARS-CoV-2 spike S1 protein

**DOI:** 10.1101/2021.01.19.427256

**Authors:** Yen-Hung Lin, Yang Han, Abhinav Sharma, Wejdan S. AlGhamdi, Chien-Hao Liu, Tzu-Hsuan Chang, Xi-Wen Xiao, Akmaral Seitkhan, Alexander D. Mottram, Pichaya Pattanasattayavong, Hendrik Faber, Martin Heeney, Thomas D. Anthopoulos

**Affiliations:** Department of Physics, Imperial College London, London SW7 2AZ, U.K.; Clarendon Laboratory, Department of Physics, University of Oxford, Oxford OX1 3PU, U.K.; Department of Chemistry, Imperial College London, London SW7 2AZ, U.K.; School of Materials Science and Engineering, Tianjin University, Tianjin 300072, P.R. China; King Abdullah University of Science and Technology (KAUST), KAUST Solar Centre, Thuwal 23955-6900, Saudi Arabia; Department of Mechanical Engineering, National Taiwan University, Taipei 10617, Taiwan; Department of Electrical Engineering, National Taiwan University, Taipei 10617, Taiwan; Department of Materials Science and Engineering, School of Molecular Science and Engineering, Vidyasirimedhi Institute of Science and Technology (VISTEC), Rayong 21210, Thailand

**Author notes:** These authors contributed equally to this work (Yen-Hung Lin, Yang Han).

## Abstract

Solid-state transistor sensors that can detect biomolecules in real time are highly attractive for emerging bioanalytical applications. However, combining cost-effective manufacturing with high sensitivity, specificity and fast sensing response, remains challenging. Here we develop low-temperature solution-processed In_2_O_3_/ZnO heterojunction transistors featuring a geometrically engineered tri-channel architecture for rapid real-time detection of different biomolecules. The sensor combines a high electron mobility channel, attributed to the quasi-two-dimensional electron gas (q2DEG) at the buried In_2_O_3_/ZnO heterointerface, in close proximity to a sensing surface featuring tethered analyte receptors. The unusual tri-channel design enables strong coupling between the buried q2DEG and the minute electronic perturbations occurring during receptor-analyte interactions allowing for robust, real-time detection of biomolecules down to attomolar (aM) concentrations. By functionalizing the tri-channel surface with SARS-CoV-2 (Severe Acute Respiratory Syndrome Coronavirus 2) antibody receptors, we demonstrate real-time detection of the SARS-CoV-2 spike S1 protein down to attomolar concentrations in under two minutes.

## Main text

Miniaturised biochemical sensors fabricated via high-throughput manufacturing methods promise cost-effective, large-volume production for use in various technology sectors^1^. The present needs for biochemical detection are diverse and include environmental monitoring^2^, security systems^3^, and preventative medical care^4^. An ideal biochemical sensing platform should be able to accommodate a wide range of applications in biological and chemical detections with high-sensitivity^5^ and selectivity^6^. Among the various types of sensing platforms, a solid-state transistor sensor is a highly-anticipated tool that could address these requirements as it provides the functionality of a transducer for converting a biochemical interaction into an amplified electrical signal^7^. This characteristic enables direct readout without the need of bulky peripheral driving (opto)electronics, such as amplifiers, excitation light sources and photo-detectors^8^.

For the successful use of solid-state transistors as biosensors, the channel should exhibit a large surface area^9^ and tuneable surface chemistry^10^. The former allows tethering of a sufficient quantity of molecular receptors whilst the latter helps to preserve charge transport in the channel without unintentionally reacting with the environment. One widely reported biosensor technology platform is based on silicon-nanowire (Si-NW) transistors, but their manufacturing remains technologically demanding^11,12,13^. Alternative technologies such as solid-state thin-film transistors (TFT) made of metal oxide semiconductors offer scalable manufacturing and intriguing physical properties^14-17^. However, due to parasitic gating effects and associated performance deterioration^18-21^, the use of metal oxide transistors as biosensors has remained limited with most effort dedicated on liquid-gated transistors (LGTs)^6,22-24^. In spite of being one of the most studied device, LGT biosensors face the detrimental Debye screening effect^6,25,26^ – a direct result of the operating principles that rely on electrochemical reactions^27^, or on the movement of analytes^28^, upon liquid-gating. Thus managing or overcoming the Debye screening effect is critical for developing ultra-sensitive transistor-based sensor technologies for emerging applications^29^.

Here, we introduce a nanometres-thin In_2_O_3_/ZnO heterojunction channel and combine it with a geometrically engineered tri-channel architecture several millimetres in size as a universal platform for rapid, selective and ultra-sensitive biosensing. The all-solid-state device features a central sensing channel and two side channels featuring a quasi-two-dimensional electron gas (q2DEG) formed at the buried heterointerface a few nanometres below the channel’s surface. The flexible surface chemistry of metal oxides, on the other hand, allows direct functionalisation of different receptors. The unusual channel architecture offers ultrahigh surface-area-to-volume ratio (10^6^ cm^2^ cm^-3^) and facilitates nm-proximity between the electrostatic perturbations occurring during surface-tethered receptor and analyte interaction, with the buried q2DEG channel. These unique features enable simultaneous signal transduction and amplification in an all-solid-state TFT platform enabling real-time detection of specific biomolecules down to attomolar (aM) concentrations under physiological relevant conditions. As a proof-of-concept we demonstrate selective sensing of the SARS-CoV-2 (Severe Acute Respiratory Syndrome Coronavirus 2) spike S1 protein in real-time with a limit of detection (LoD) of 865 aM.

### Quasi-two-dimensional oxide heterojunction channel

We hypothesised that our recently developed solution-processed, high electron mobility In_2_O_3_/ZnO heterojunction (HJ) transistors^30^ offers unique features that could prove attractive for biosensing. Firstly, the buried electron channel located at the oxide HJ is physically separated from the receptor units tethered on its surface a few nm above^31,32^. This feature is expected to prevent degradation of electron transport upon sensing (due to Coulomb scattering) and preserve the transistor’s performance. This is not the case for most biosensor transistors reported to date where the channel interacts directly with the receptor units and hence the analyte. To overcome this, liquid gating has been exploited for analyte detection in the liquid phase^6,22,33^. Secondly, the high electron mobility of the HJ TFTs offers the possibility for large electrical signals that are easy to detect and amplify even in large-size devices^31^.

We fabricated metal oxide HJ transistors using the staggered bottom-gate, top-contact (BG-TC) architecture shown in **Fig. 1a**. High-resolution transmission electron microscopy (HRTEM) analysis (**Fig. 1b)** of the channel reveals the formation of a well-defined HJ channel with thickness in the range of 8-10 nm. Atomic force microscopy (AFM) measurements show the existence of smooth layers as being deposited sequentially (**Fig. 1c-d)**. In_2_O_3_ exhibits the lowest peak-to-peak height (ΔZ) of 1.87 nm with a root-mean-square roughness (σ_RMS_) value of 0.20 nm, which are comparable to that of SiO_2_ (ΔZ = 1.91 nm, σ_RMS_ = 0.21 nm). Subsequent deposition of ZnO atop In_2_O_3_ leads to a slightly rougher topography (ΔZ = 4.00 nm, σ_RMS_ = 0.58 nm) indicative of a more textured surface^31,34^.

**Figure 1.**
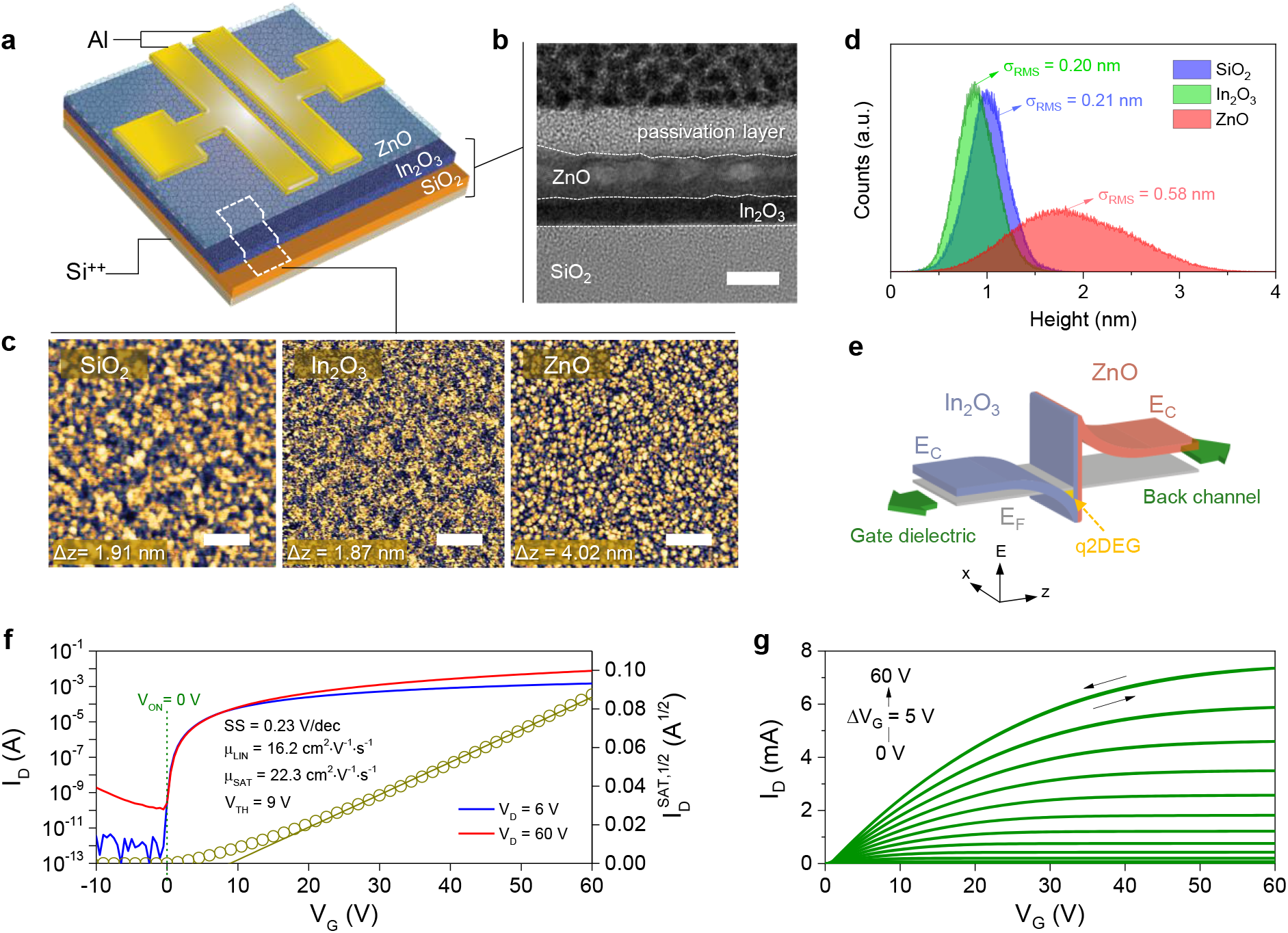
Fabrication and testing of metal oxide heterojunction transistors. **a**, Schematic of an In_2_O_3_/ZnO heterojunction transistor. **b**, HRTEM cross-sectional image of the channel region (scale bar = 5 nm). **c**, Intermittent AFM topography images of SiO_2_, In_2_O_3_ and ZnO surfaces (scale bar = 200 nm). (**d**) Height histogram extracted from the AFM data for each sequentially deposited layer. Corresponding peak-to-peak height difference (ΔZ) and root mean square surface roughness (σ_RMS_) were derived from AFM image analysis. **e**, Schematic of energetic diagram for the In_2_O_3_/ZnO heterointerface. The discontinuity in the conduction band between ZnO and In_2_O_3_ results to the electron migration from ZnO to In_2_O_3_, resulting in the formation of a q2DEG. **f**,**g**, Representative current-voltage (I-V) characteristics for a In_2_O_3_/ZnO transistor: (**f**) transfer and (**g**) output characteristics. Important device parameters are shown in (**f**). These include turn on voltage (V_ON_), threshold voltage (V_TH_), subthreshold swing (SS), linear mobility (µ_LIN_), and saturation mobility (µ_SAT_).

The In_2_O_3_/ZnO forms a type-II heterojunction where electrons migrate from the conduction band (CB) of ZnO to that of In_2_O_3_, leading to the formation of a q2DEG (**Fig. 1e**)^32^. The latter resembles 2DEG systems found in high electron mobility transistors (HEMTs) based on epitaxial inorganic heterointerfaces^35^. Although uncommon, the existence of q2DEG in disordered/non-epitaxial heterointerfaces has been predicted, further corroborating our findings and conclusions^36^. **Fig. 1f** and **1g** show representative sets of transfer and output current-voltage (I-V) characteristics for a In_2_O_3_/ZnO HJ transistor with electron mobility and current on-off ratio of >22 cm^2^ V^-1^ s^-1^ and >10^8^, respectively. The presence of the q2DEG system at the In_2_O_3_/ZnO interface is responsible for the electron current across the channel and the high electron mobility measured^32^.

### All solid-state tri-channel transistor sensor

To investigate the suitability of the In_2_O_3_/ZnO transistors for biosensing, we fabricated devices based on a tri-channel configuration on 4-inch Si wafers (**Fig. 2a**). The source-drain (S-D) electrodes are deposited atop the In_2_O_3_/ZnO channel followed by the deposition of another ultrathin (2-4 nm) protective ZnO layer. Next the known deoxyribonucleic acid (DNA) intercalator^37,38^ 1-pyrenebutyric acid (PBA) was functionalised directly to ZnO^39^ acting as the DNA receptor. A second functionalisation step using butyric acid (BA) was also applied to ensure complete passivation of the ZnO surface (see **Supplementary Note 1** and **Supplementary Figure 1**). The presence of the PBA molecules was verified using ultraviolet– visible (UV-Vis) absorption measurements before and after functionalisation as evidenced by the appearance of distinct absorption peaks associated with the pyrene unit (**Supplementary Figure 2**). The completed device consists of two identical ‘conventional’ channels (hereafter termed CC) 100 μm in length (L), formed on the sides, and a third long (L = 2000 μm) ‘sensing’ channel (hereafter termed SC) formed in the central region of the device between the S-D electrodes (see **Supplementary Figure 3**). This unique channel layout offers a large sensing area channel where the analyte-containing solution can be easily applied while avoiding direct contact with the S-D electrodes (**Fig. 2b-c**) which is known to induce parasitic gating effects^40^.

**Figure 2.**
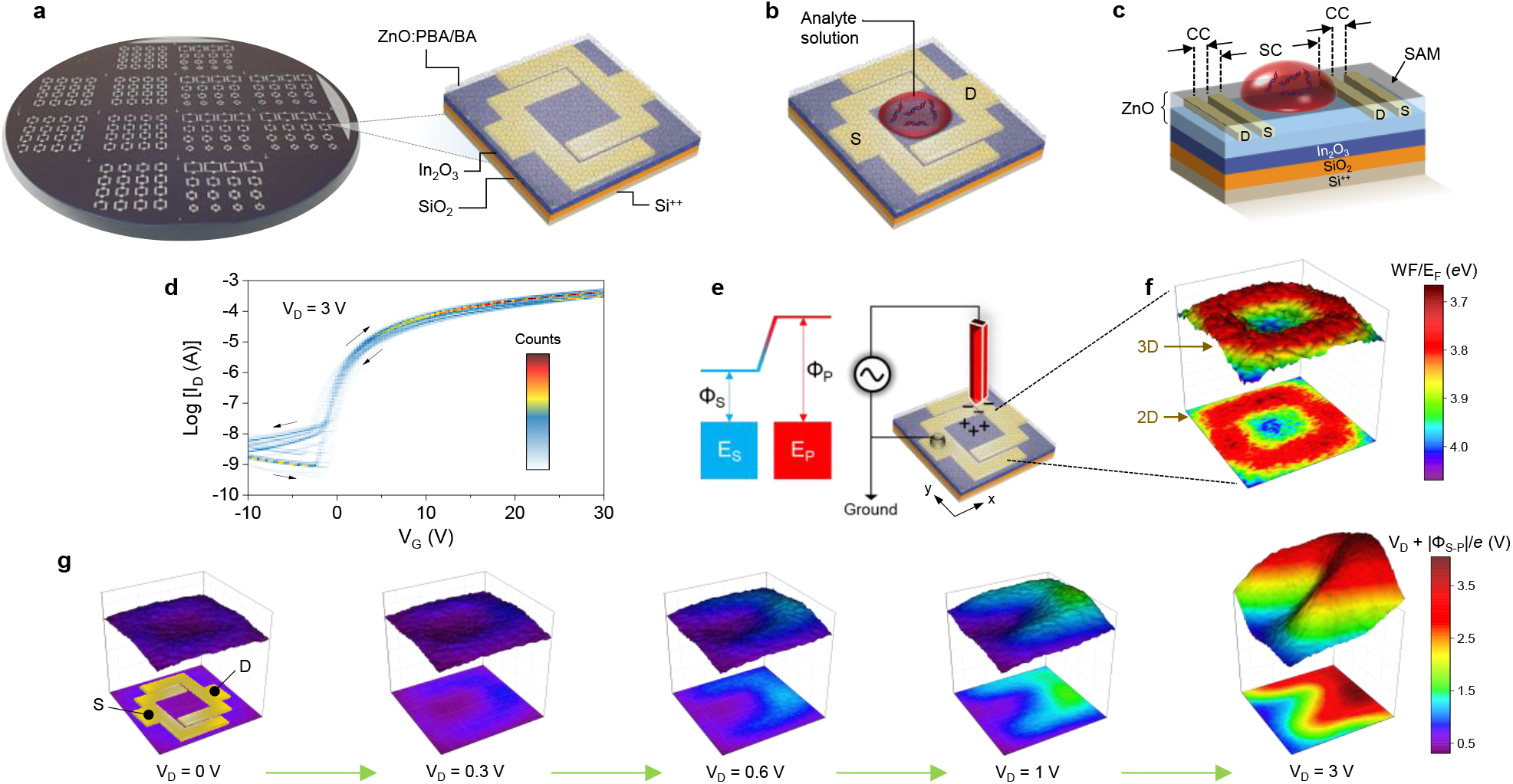
Design and structures of tri-channel transistor sensors. **a**, Tri-channel In_2_O_3_/ZnO heterojunction transistors fabricated on a 4-inch Si^++^/SiO_2_ wafer and schematic of the channel architecture. The source-drain (S-D) electrodes are covered by the top ZnO layer. The receptor molecule pyrenebutyric acid (PBA) and passivation molecule butyric acid (BA) are chemically tethered onto the ZnO surface. **b**, Illustration of the direct application of analyte solution on the millimetre-scale sensing channel (SC) area of the sensor. **c**, Schematic of the tri-channel transistor depicting the location of the analyte solution within the SC and two conventional channels (CCs) on the sides. **d**, Density plots of forward-backward dual sweeps of current-voltage characteristics measured from 30 individual tri-channel transistor sensors. **e**, Schematic of the scanning Kelvin probe (SKP) setup used. The SKP method relies on the application of a voltage to offset the surface potential between the sample (σ_S_) and the tip (σ_P_). The magnitude of this voltage is then used to calculate the energy difference between the sample (E_S_) and the tip (E_P_). **f**, 2D (top)/3D (bottom) maps of the electrostatic potential across a tri-channel transistor measured by SKP. The WF for the embedded Al-electrode areas is measured to be ≈ 3.8 eV while the E_F_ for the SC is ≈ 4.0 eV. **g**, Electrostatic potential maps measured at different source-drain potentials: V_D_ = 0, 0.3, 0.6, 1, 3 V. The relative positions of the S-D electrodes are shown in the 2D map for V_D_ = 0 V.

In **Supplementary Figures 4** and **5**, we plot the transistor transfer and output characteristics, respectively, measured before and after PBA and BA functionalisation. Unlike conventional transistor biosensors^22,41^, our tri-channel device shows negligible changes in its operating characteristics after receptor functionalisation. The narrow parameter distribution is better illustrated in **Fig. 2d** which shows the density plots^42^ of the dual-sweep transfer characteristics for 30 individual tri-channel transistors fabricated on a single wafer. Critically, the tri-channel transistors exhibit robust operation even when subjected to 90 repeated dual I-V sweeps with negligible leakage current (I_G_) which is critical for optimal device operation and signal amplification (**Supplementary Figure 6)**^43^. These data demonstrate the high operational stability and reproducibility of the proposed tri-channel HJ transistor architecture.

To better understand the electrostatic potential landscape across our unconventional tri-channel device, we performed scanning Kelvin probe (SKP) measurements (**Fig. 2e**). **Fig. 2f** shows the two-dimensional (2D, bottom) and three-dimensional (3D, top) work function (WF) or Fermi energy (E_F_) maps for a tri-channel device measured. The influence of the buried Al electrodes beneath the ZnO results in local WF changes (3.8-4 eV), with the higher potential observed in the middle of the SC region. SKP measurements were also performed while applying a drain bias (V_D_) in the range of 0-3 V (**Fig. 2g**; the respective location of the device illustrated for V_D_ = 0 V image). The application of low voltages (e.g. V_D_ = 0.6-1 V) causes a substantial change within the SC, while increasing the applied bias to 3 V affects the potential landscape across the entire SC region, suggesting strong coupling between the SC and the two side CCs. Thus, the tri-channel architecture appears to enable spatially decoupling of the signal transduction occurring within the SC region from the current-driving CCs.

To understand how the tri-channel geometric impacts the electrical characteristics of the sensor, we modelled the device operation using the COMSOL Multiphysics^®^ simulation software. **Fig. 3a** shows the simulated and measured transfer characteristics for a representative transistor. The various material and device parameters used in the modelling were adopted from our previous studies on the same materials^31,44,45^. The small difference seen in the subthreshold region between the modelled and experimental data is attributed to the presence of trap states in the channel^46,47^, while the higher off current measured experimentally is due to the use of a common Si^++^ gate substrate and the un-patterned layout of the In_2_O_3_/ZnO channel. Apart from these minor discrepancies, the model provides a good description of the tri-channel transistor operation and validates its applicability.

**Figure 3.**
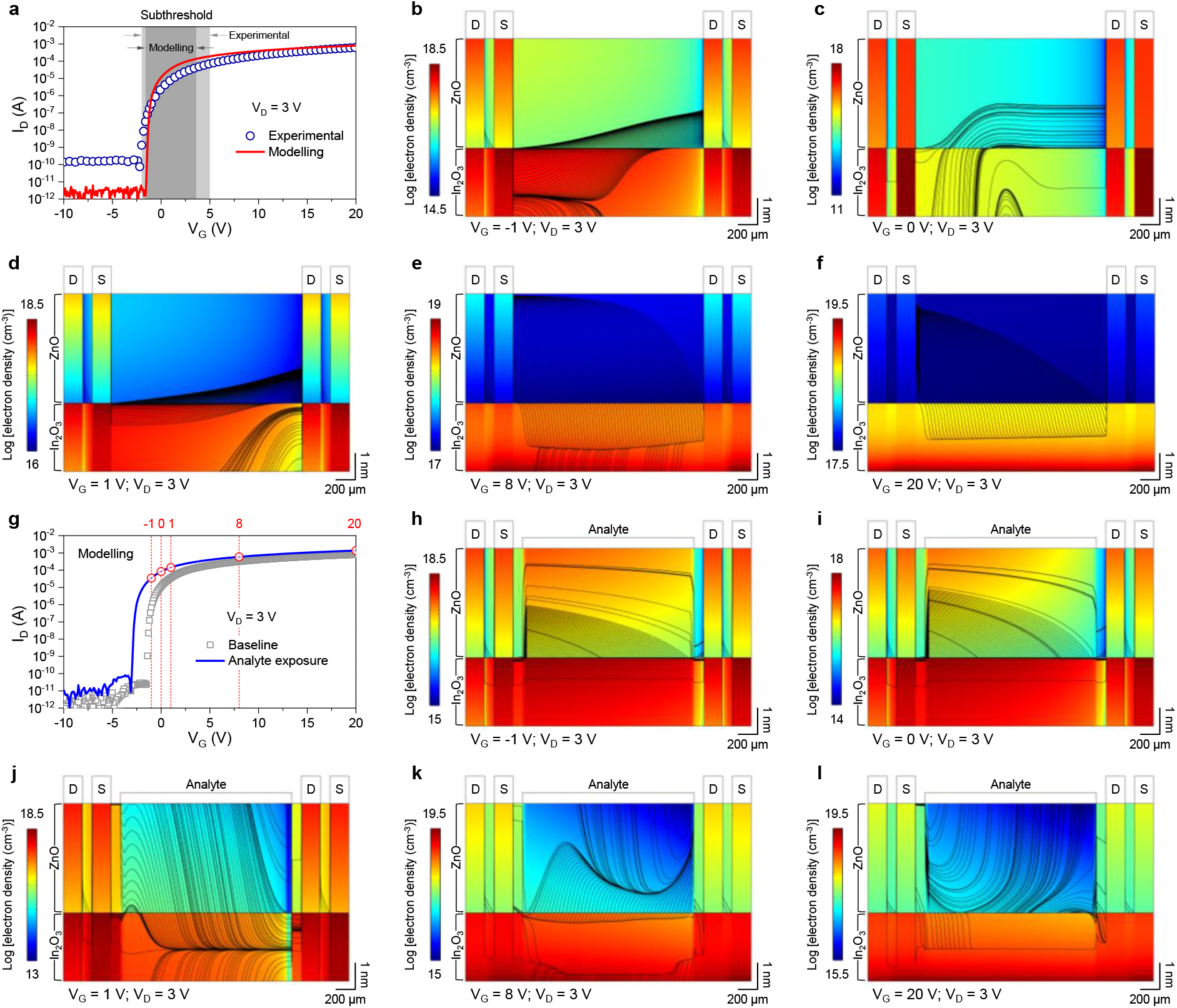
Physical principles of tri-channel transistor sensors. **a**, Transfer current-voltage characteristics of tri-channel transistor sensors obtained from experiment and modelling using COMSOL Multiphysics^®^. The applied drain voltage (V_D_) was +3 V, and the subthreshold regions are indicated in grey. **b**,**c**,**d**,**e**,**f**, Corresponding COMSOL simulations showing the electron density distributions along the cross-section of the In_2_O_3_/ZnO heterostructure under the source (S) and drain (D) electrodes and the electron flow streamlines within the channel regions, with different gate voltages (V_G_) applied: (**b**)-1 V; (**c**) 0 V; (**d**) 1 V; (**e**) 8 V; (**f**) 20 V, and a constant V_D_ = 3 V. **g**, Modelled transfer current-voltage characteristics (V_D_ = 3 V) for baseline and under the exposure of simulated surface-charged analytes. **h**,**i**,**j**,**k**,**l**, Corresponding COMSOL electron density distributions under the influence of simulated analytes when applying V_D_ = 3 V and V_G_ = (**h**)-1 V; (**i**) 0 V; (**j**) 1 V; (**k**) 8 V; (**l**) 20 V to the sensors. The electrodes and analytes are shown to indicate their positions with respect to the devices.

The main function of a transistor biosensor is to induce a perturbation in the channel current upon exposure to an external stimulus (analyte). To best illustrate this process in our sensor, we used the post-processing streamline tool for visualising the electron concentration and the streamlines of the channel current flow. **Fig. 3b-f** show the static distributions of the electron density and the streamlines of the current flow within the In_2_O_3_/ZnO heterostructure biased at V_D_ = 3 V and V_G_ =-1, 0, 1, 8 and 20 V. The results are in good agreement with our experimental observations and reveal the staggering enhancement in the current density within the In_2_O_3_ of the heterointerface^31,46^. Next, we modelled the electrical characteristics of the device in the presence of an analyte. We hypothesize that the analyte species interacts with the surface-tethered receptor units and induce free charges at the surface of the SC region. To establish the sensing condition close to the limit of detection for our sensor, we assumed the number of additional charges induced by the analyte to be equivalent or lower than the number of mobile charges in the channel. Based on the literature^48^ and our own measurements on similar metal oxide heterointerface channels^31^, device operation should remain largely unaltered when the additional electron concentration remains below 10^17^ cm^-3^.^34^ To ensure that this condition is satisfied, we used a more conservative estimation for the analyte-induced electron concentration of 10^16^ cm^-3^ and a channel thickness of 10 nm (**Fig. 1b**). The equivalent surface charge density due to analyte was then derived from the modelling yielding a value of ≈10^10^ cm^-2^, which will be considered for the different device operating scenarios next.

**Fig. 3g** shows the modelled transfer characteristics for a tri-channel (inset, **Fig. 3g**) In_2_O_3_/ZnO sensor while **Supplementary Figure 7** displays similar calculations for a single layer In_2_O_3_ and a heterojunction In_2_O_3_/ZnO transistors based on the conventional channel geometry (insets in **Supplementary Figure 7a**,**c**) before (baseline) and after exposure to analyte (analyte exposure). The tri-channel In_2_O_3_/ZnO transistor shows a large response to the analyte (surface charge ≈10^10^ cm^-2^) with the transfer curve shifted towards more negative V_G_ bias. This is not the case for In_2_O_3_ and In_2_O_3_/ZnO transistors with conventional channel geometry (**Supplementary Figure 7b**,**d**), where the analyte induces only a small perturbation in the current around the subthreshold region consistent with filling of sub-gap states^49^. The modelled electron density and current flow for the tri-channel transistor biased at V_D_ = 3 V and V_G_ = −1, 0, 1, 8 and 20 V, are presented in **Fig. 3h-l**, whilst the corresponding modelling results for the conventional channel In_2_O_3_ (at V_D_ = 3 V, V_G_ = 1 V) and In_2_O_3_/ZnO (at V_D_ = 3 V, V_G_ = −1 and 1 V) transistors are shown in **Supplementary Figure 8a-b** and **8c-f**, respectively. Strikingly, we find that unlike the geometrically engineered heterojunction In_2_O_3_/ZnO transistors (**Fig. 3h-l**), electron flow in the In_2_O_3_ device is pinned at the interface with the gate dielectric while being fully decoupled from the surface/analyte (**Supplementary Figure 8b**). From these data we conclude that the tri-channel design is highly sensitive to the presence of surface charges as compared to conventional channel design, while single layer In_2_O_3_ channel transistors are not ideal for all-solid-state biosensing applications.

Next, we considered the scenario where the heterojunction transistor is operated in depletion (V_G_ = −1 V) and in the presence of analyte (i.e. additional ≈10^10^ cm^-2^ on the SC surface). Clear perturbations in the current flow are observed for both the tri-channel In_2_O_3_/ZnO (**Fig. 3h**-**l**) and the conventional channel designs of In_2_O_3_/ZnO (**Supplementary Figures 8d** and **8f**) transistors. The broader distribution of streamlines seen in the tri-channel is consistent with the large negative shift in the turn-on voltage (V_ON_) of the device seen in **Fig. 3g**. Regardless of the biasing scenarios (depletion or accumulation), the tri-channel architecture shows much stronger coupling to the analyte. Specifically, we find the electron flow streamlines to extend ≈1 nm beneath the SC surface (**Fig. 3h-l**) due to the asymmetric design of the source-drain electrodes^50^, which prevent the local electric field to fully pinch-off the channel for V_G_ between-1 to 1 V. As the V_G_ increases (+20 V), the benefits associated with the presence of a q2DEG in the In_2_O_3_/ZnO become even more apparent as the area beneath the sensing surface remains free from electrostatic screening induced by the gate (**Fig. 3l**). Nevertheless, it is known to be more advantageous for solid-state transistor sensors to be operated within the subthreshold region as it yields optimal sensitivity due to high signal gain^51^.

### Receptor engineering for ultra-sensitive and real-time biosensing

To demonstrate that the working principle of our all-solid-state tri-channel transistor is fundamentally different from that of conventional liquid-gated sensors, we studied the ability of our transistors to detect different types of DNAs (analytes) dispersed in deionised (DI) water rather than in a high ionic strength solution. We note that the latter is essential for the function of liquid-gated transistor sensors in order to drive the analyte towards the semiconducting channel, which in turn modulates its transconductance via electrochemical processes^6^. To prove that our sensors do not rely on such processes, the DI-water based solutions containing double-stranded DNA (dsDNA) and single-stranded DNA (ssDNA) of different sequences, were applied directly onto the SC area while recording the device’s response. **Fig. 4a** depicts the envisioned interaction between dsDNA and PBA where the pyrene units on PBA intercalate into the dsDNA^52^. **Fig. 4b-d** show the measured transfer characteristics (V_D_ = 3 V) for different concentrations of 20 base-pair segments of synthetic DNAs based on single-stranded adenine [abbreviated as A20], and thymine (T) [abbreviated as T20], as well as their complementary dsDNA (AT)20. For (AT)20, a much larger change in the transistor’s transfer characteristics is observed with the lowest dsDNA concentrations studied down to 100 aM (**Fig. 4b**). The strong response is attributed exclusively to the intercalation of the pyrene units into the minor grooves of the double-stranded (AT)20 since the presence of DI water has no measurable effect (**Supplementary Figure 9**). The progressive shift of V_ON_ towards more negative V_G_ seen in **Fig. 4b** is consistent with the modelling results of **Fig. 3g** where we considered the presence of additional free charges on the surface of the SC. This observation indicates that pyrene-NDA association generates free electrons that are eventually injected into the channel. Further evidence supporting our hypothesis comes from sensing experiments involving the single-stranded A20 (**Fig. 4c**) and T20 (**Fig. 4d**) where only minute changes are observed in the transistors’ characteristics due to the absence of pyrene-NDA intercalation.

**Figure 4.**
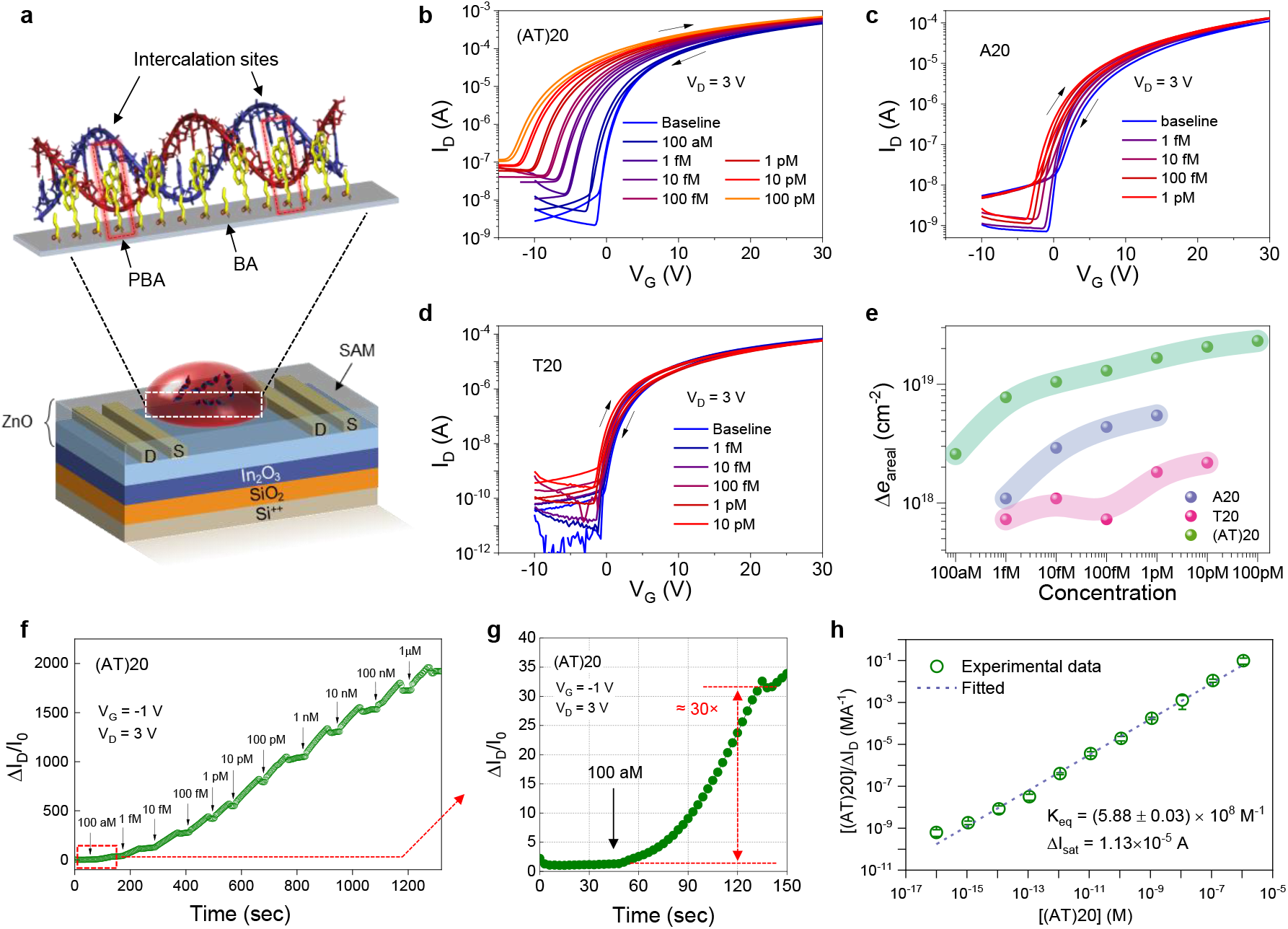
Tri-channel transistor sensor for synthetic DNA sensing. **a**, Illustration of the envisioned intercalation between the pyrene units and dsDNA. **b**,**c**,**d**, Transfer I-V characteristics (V_D_ = 3 V) measured from PBA/BA functionalised tri-channel transistor sensors with the presence of three different DNA analytes of (**b**) (AT)20; (**c**) A20; (**d**) T20 at different analyte concentrations. **e**, Plot of the increase in areal charge carriers Δ*e*_areal_ that results from the sensing activity of the tri-channel transistor sensor to the analytes as a function of analyte concentration. Δ*e*_areal_ is calculated from the shift in the turn-on voltage of the device upon the application of analyte solution. AT(20) shows the highest response due to its interaction via intercalation with pyrene units of the PBA-functionalised tri-channel transistor sensor. **f**, Real-time response signal measured from a PBA/BA functionalised solid-state tri-channel transistor sensor operated at V_G_ =-1 V and V_D_ = 3 V upon exposure to synthetic (AT)20 with concentrations from 100 aM to 1 µM. **g**, recorded response to 100 aM showing ≈30 times enhancement in I_D_. The arrows indicate the time when the different analyte concentrations were applied to the SC area of the tri-channel transistor. **h**, Fitting of experimental results of synthetic AT(20) sensing at different analyte concentrations according to the Langmuir adsorption isotherm. The error bars denote standard deviations from three real-time measurement sets.

In an effort to quantify the sensor’s response, we analysed the change in V_ON_ as a function of increasing analyte concentration. This shift reflects the increase in the electron concentration (Δ*e*_areal_) within the channel and is given as^53^

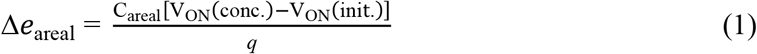

Here, C_areal_ is the areal capacitance of the gate dielectric (34.4 nF cm^-2^), *q*is the elementary charge, V_ON_(init.) is the initial V_ON_ measured in the presence of blank solution (no analyte), and V_ON_(conc.) is the transistor’s V_ON_ measured upon application of the analyte at each concentration. For simplicity, we assume all electrons are confined in a two-dimensional plane at the vicinity of the oxide HJ^32^. **Fig. 4e** shows the evolution of Δ*e*_areal_ as a function of analyte concentration measured using a tri-channel sensor. (AT)20 induces the highest Δ*e*_areal_, a direct consequence of the large V_ON_ shift observed in **Fig. 4b**. These results demonstrate unambiguously that pyrene-(AT)20 intercalation produces signals several orders of magnitude larger than the non-intercalating ssDNAs A20 and T20 and showcase the ability of the tri-channel sensor to differentiate between double and single stranded DNAs without the need for complex fluorescence labelling^54^. To this end, the DNA conformation with respect to the substrate (i.e. lying-down or standing up), should not be critical as sensing relies exclusively on the charge transfer upon pyrene-DNA association. This hypothesis is corroborated by the sensor’s ability to selectively detect different analytes, such as avidin and SARS-CoV-2 spike S1 protein, which will be discussed latter. The ability of our sensor to facilitate such a strong coupling between the minute receptor-analyte interactions and charge transport, without compromising the channel transconductance (g_m_) (**Supplementary Note 2** and **Supplementary Figure 10**), is attributed to three unique device attributes:

i. The geometrical engineered tri-channel design that enables strong coupling between current transport and receptor-analyte interactions within the sensing area of the channel.
ii. The use of a high electron mobility In_2_O_3_/ZnO channel featuring a spatially-separated (buried) q2DEG system.
iii. The versatile surface chemistry and the electronic properties of the metal oxide employed.

Due to the diverse range of biosensor transistor technologies^5,6^ there is currently no clear consensus on the important figures of merit that can be used to define the performance of such devices. Here we attempt to draw an analogy from the field of phototransistors, since both types of sensors act as transducers with a highly V_G_-dependent response and define two practical figures of merit namely the responsivity (R_analyte_) and sensitivity (S_analyte_) (**Supplementary Note 3** and **Supplementary Figure 11**). We first investigated the suitability of our tri-channel biosensor TFTs for real-time sensing of (AT)20 at an extremely broad range of analyte concentrations (10^−18^ to 10^−6^ M), while simultaneously assessing the sensors’ ability to operate in aqueous conditions^9,55^. Specifically, we monitored the evolution of ΔI_D_ at V_G_ = −1 V and V_D_ = 3 V, as a function of time for different (AT)20 concentrations. The biasing condition were chosen to maximise the sensor’s response by operating it in the subthreshold region^51^ (**Supplementary Figure 12**). **Fig. 4f** shows a representative real-time recording of ΔI_D_/I_0_ (where I_0_ = 3.16×10^−8^ A) for analyte concentrations in the range 10^−18^ to 10^−6^ M, where a clear response across the entire range is observed. Even at 100 aM of (AT)20, the tri-channel TFT shows a significant increase in the ΔI_D_ by ≈ 30 times (**Fig. 4g**) in under 2 min. This represents the highest response signal reported to date for biosensing transistors, including liquid-gated devices^5,6,24^. Importantly, the sensor’s sensitivity can be tuned by the V_G_ as shown in **Supplementary Figure 13** where the S_analyte_ is plotted *vs*. (AT)20 concentration for different V_G_ (−1, 0, +1 V). Even at sub-optimal biasing conditions (i.e. V_G_ = 1 V), the measured I_D_ for 100 aM (AT)20 increases to 4.2 µA (ΔI_D_ ≈ 2.8 µA) which is ≈ 300% higher than the baseline signal (I_0_ ≈ 1.4 µA) (**Fig. 4b**). The large ΔI_D_ indicates that the actual sensitivity of the tri-channel sensors is well below 100 aM. To this end, we note that in the literature the most frequently reported parameter is the LoD, which is determined by the minimum detectable signals that are often far from suitable for real-time monitoring ^5,6,41,56^.

To further demonstrate the capabilities of our high S_analyte_ sensor, we analysed the sensing kinetics using the linear form of the Langmuir adsorption isotherm^57,58^. **Supplementary Figure 14a** displays a series of such measurements taken from **Fig. 4f** but replotted by setting the time (t) at which the different concentrations of analyte were applied, to 0 sec. **Supplementary Figure 14b** shows a representative trace for 1 pM of (AT)20 where three different sensing stages can be clearly distinguished:

i. Concentration-limited diffusion stage where the receptor-analyte reaction rate is determined by the diffusive transport of the analyte on the sensor’s surface as its concentration increases.
ii. Association of analyte with the tethered receptor moieties i.e. ‘primary’ sensing process.
iii. Dissociation of analyte-receptor complexes before reaching a thermodynamic equilibrium.

The rate of the ‘primary’ sensing process depicted in **Supplementary Figure 14b** is representative of a zero-order reaction and is independent of the analyte concentration or the method with which the analyte solution is being applied. For each concentration, a distinct peak between association and dissociation stages is observed and attributed to the immobilisation of analyte species by the tethered receptors^59-61^. Therefore, and regardless of the sensing method, the existence of two-phase kinetics relates solely to the association and dissociation stages. Using the high fidelity sensing data from **Supplementary Figure 14**, the equilibrium constant (K_eq_) was calculated yielding values of (5.88 ± 0.03) × 10^8^ M^-1^ (**Fig. 4h**).

In addition to short synthetic DNA, we have also tested natural dsDNA extracted from calf thymus tissue, which has much longer DNA sequences. **Fig. 5a**-**b**, respectively, show the transfer characteristics (V_D_ = 3 V) and real-time response recorded at fixed V_D_ = 3 V and V_G_ = −1 V (**Supplementary Figure 15**, I_0_ = 2.63 × 10^−8^ A). The response is similar to that recorded for (AT)20 indicating that the sensing mechanism remains identical for the natural dsDNA. Even when aM concentration of the dsDNA is applied, the recorded signal (ΔI_D_/I_0_) increases by more than 100× (inset of **Fig. 5b**), further corroborating the unprecedented sensitivity of the tri-channel sensor. When compared to (AT)20, the sensor exhibits stronger response to natural dsDNA with a higher binding constant K_eq_ of (8.71 ± 0.01) ×10^9^ M^-1^ (**Fig. 5c**). This difference is attributed to the stronger interaction between the longer sequence of calf thymus DNA and the surface-tethered pyrene receptor.

**Figure 5.**
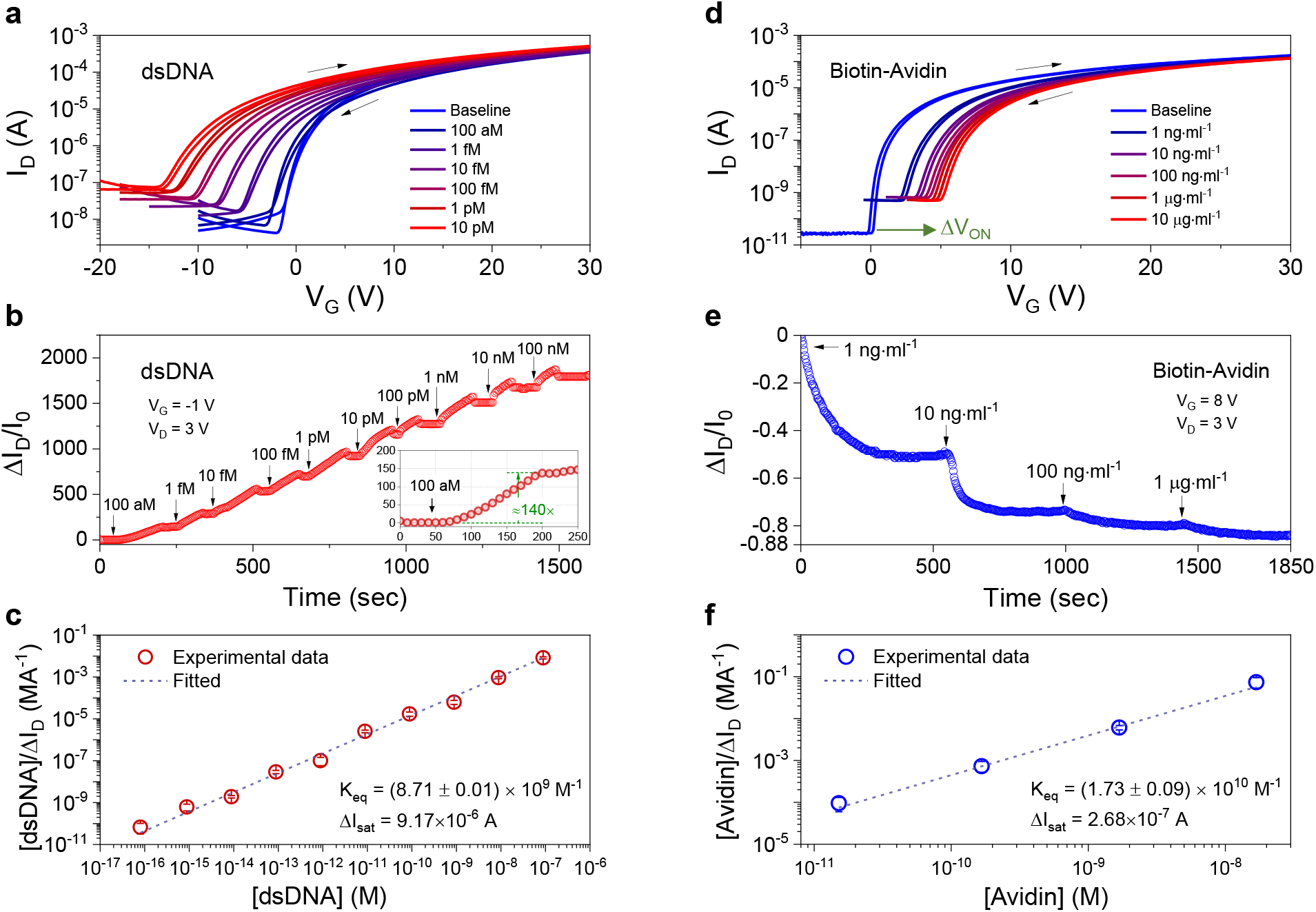
Attomolar detection of natural biomolecules. **a**, Transfer characteristics (V_D_ = 3 V) of a PBA/BA functionalised tri-channel transistor sensor measured in the presence of natural dsDNA extracted from calf thymus. **b**, Real-time response of the tri-channel transistor sensor to different concentrations (100 aM to 100 nM) of natural dsDNA. Inset: The sensor’s response to a 100 aM of the analyte is ≈140 times higher than the baseline signal. For this experiment the device was operated at V_G_ =-1 V and V_D_ = 3 V. **c**, Fitting of the experimental results for natural dsDNA at different analyte concentrations according to the Langmuir adsorption isotherm. The error bars denote standard deviations from three real-time measurement sets. **d**, Transfer characteristics (V_D_ = 3 V) measured from a biotin-functionalised tri-channel transistor sensor subject to different concentrations of avidin. **e**, Real-time response obtained from the biotin-based tri-channel transistor sensor biased at V_G_ = 8 V and V_D_ = 3 V. The avidin concentration was varied from 10 ng ml^-1^ to 1 µg ml^-1^. The arrows indicate the time when the avidin was applied to the SC area of the sensor. **f**, Fitting of experimental results of avidin sensing at different analyte concentrations according to the Langmuir adsorption isotherm. The error bars denote standard deviations from three real-time measurement sets.

We also investigated the possibility of sensing the formation of the positively charged biotin-avidin pair – an important complex for biochemical analysis^62^. For this purpose, we first functionalised the ZnO surface with biotin, acting as the receptor, and then applied avidin (the analyte) dispersed in DI solution at different concentrations. **Fig. 5d** reveals a systematic shift in V_ON_ of the transistors towards more positive V_G_ with increasing avidin concentration. This indicates a continuously reducing electron concentration in the channel due to the positively charged nature of avidin and its electron accepting character. **Fig. 5e** shows real-time sensing of different concentrations of avidin. Here a higher voltage bias of V_D_ = 3 V, V_G_ = 8 V (**Supplementary Figure 16**, I_0_ = 1.89 × 10^−5^ A) was used in order to compensate for the large positive shift in V_ON_ upon biotin-avidin association. Analysis of the binding constant between avidin and surface-immobilised biotin (**Fig. 5f**) yields a K_eq_ of 1.73 (±0.09) × 10^10^ M^-1^. This value is lower than that reported for free avidin-biotin pairs (10^13^–10^15^ M^-1^)^63,64^ – a result most likely attributed to the smaller number of tethered biotin receptors.

To summarize, the sensing mechanism in our all-solid-state tri-channel transistor sensors is starkly different to that of liquid-gated sensor platforms^5,6,22-28^. The sensing process is modelled by considering the generation of free charges on the SC’s surface upon receptor-analyte association (**Fig. 3g-l**) and its strong coupling to the channel current. The higher gradients in the electron flow streamlines observed towards the HJ/analyte interface and the higher electron density highlight how excess charges are introduced and transported across the device upon receptor-analyte interaction. Importantly, the sensor can be easily repurposed via receptor engineering to detect both negatively (i.e. DNAs) as well as positively charged (i.e. biotin-avidin) analytes. In the case of biotin-avidin interaction the channel current was found to reduce due to the electron accepting nature of the formed complex. Another important feature of the tri-channel sensor is the large size SC and its ability to accommodate a high density of receptors which in turn enable dynamic sensing over an extraordinary wide range of analyte concentrations (**Supplementary Figure 14**).

### Detection of SARS-CoV-2 spike S1 protein

To demonstrate the potential of the tri-channel transistors in a real-world sensing scenario, we engineered the surface of the SC by immobilizing SARS-CoV-2 antibody acceptors designed for specific binding to the SARS-CoV-2 spike S1 protein (**Fig. 6a**)^56^. The receptor-binding domain (RBD) of the spike protein is known to bind the human cell receptor angiotensin-converting enzyme 2 (ACE2), followed by subsequent viral entry. During binding the positively charged polybasic cleavage site on the spike protein binds strongly with the negatively charged human cell receptor ACE2^65^. We hypothesized that such interaction would induce electrostatic perturbations that are detectable by the tri-channel sensor. The sensing scheme is rather straightforward and could prove highly versatile for the detection of the new corona virus and other pathogens of interest.

**Figure 6.**
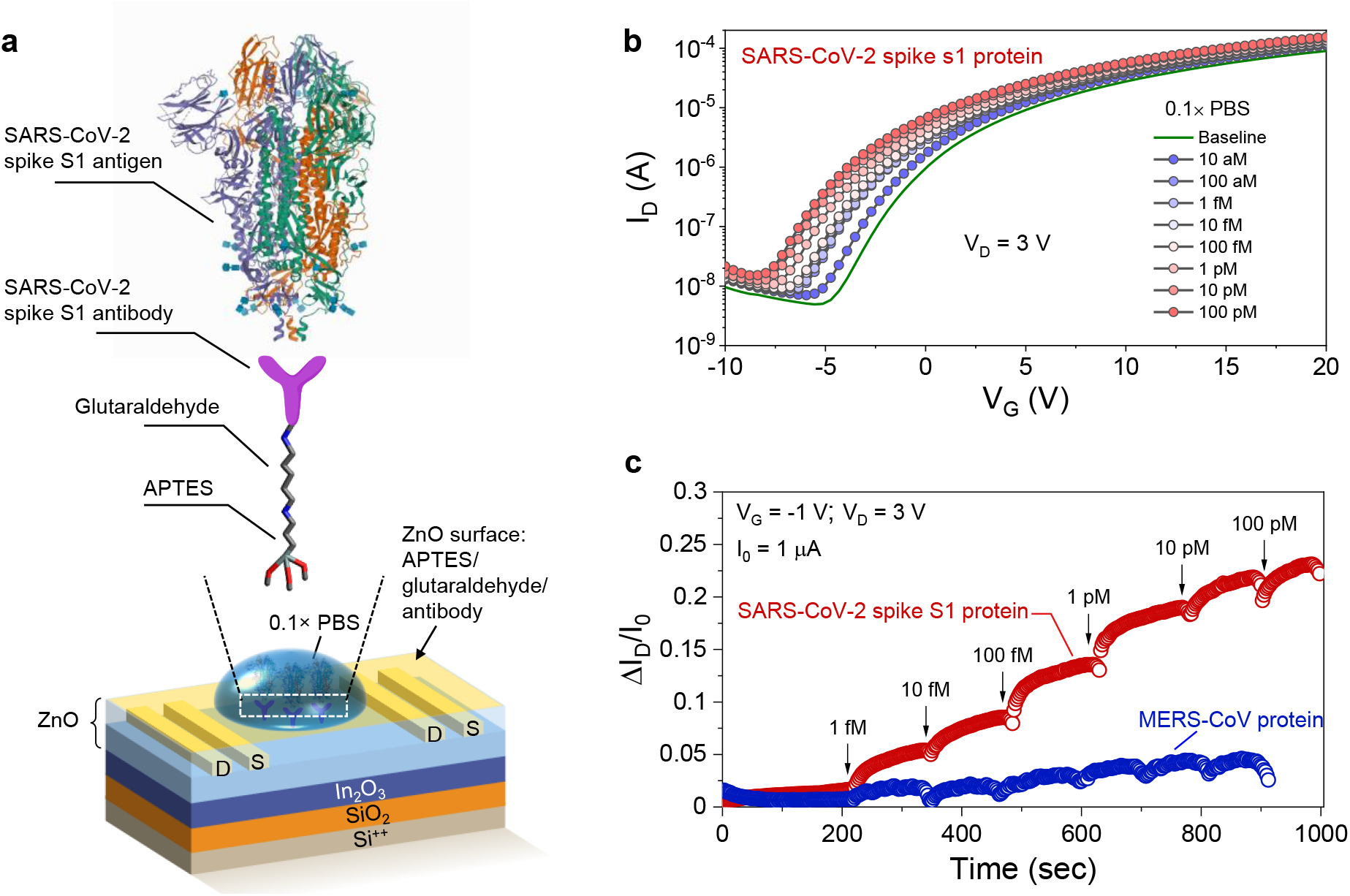
Detection of SARS-CoV-2 spike protein. **a**, Schematic of the SARS-CoV-2 spike S1 protein detection. The SARS-CoV-2 spike S1 antibody is anchored onto the sensor platform after the sequential modification of oxide surface with 3-aminopropyltriethoxysilane (APTES) and glutaraldehyde. **b**, Transfer characteristics (V_D_ = 3 V) of a fully functionalised tri-channel transistor sensor measured in the presence of the SARS-CoV-2 spike protein in 0.1× phosphate-buffered saline (PBS, baseline). **c**, Real-time response of the tri-channel transistor sensors to different concentrations (1 fM to 100 pM) of the SARS-CoV-2 spike protein and the MERS-CoV protein in 0.1× PBS.

To test our hypothesis, we first examined the ability of the sensor to operate under physiological-relevant conditions. The antibody-tethered tri-channel sensor show negligible response upon application of different concentrations of the high ionic strength phosphate buffered saline (PBS) solution onto the SC (**Supplementary Figures 17** and **18**). Next, a series of PBS solutions containing different concentrations of the SARS-CoV-2 spike S1 protein were prepared and applied to the SC of the sensor while the transfer characteristics were recorded for each analyte concentration (**Fig. 6b** and **Supplementary Figure 19a**). A clear and systematic shift in V_ON_ towards more negative gate voltages with increasing analyte concentration, is observed. Strikingly, even at 10 aM the sensor’s response remains large and clearly visible in the quasi-static transfer characteristics of **Fig. 6b** indicating a high sensitivity.

To further demonstrate the versatility of the tri-channel sensor, we performed real-time sensing measurements of the SARS-CoV-2 spike S1 protein. Prior to this the sensor was biased at V_G_ = −1 V and V_D_ = 3 V to acquire a stable baseline channel current of ≈1 µA (ΔI_D_/I_0_ = 0). Following, PBS solutions containing varying concentrations of the SARS-CoV-2 spike S1 protein where applied sequentially to the SC while the sensor current being recorded in real-time (**Supplementary Figure 19b**). Evidently, the sensor can detect the analyte across an ultra-wide range of concentrations (aM to pM) demonstrating the tremendous potential of the technology. Similar to dsDNA real-time sensing data, the recorded signal (ΔI_D_/I_0_) for each concentration increases and reaches an equilibrium followed by a small deep due to the diffusion limited, association and dissociation stages discussed previously (**Supplementary Figure 14b**).

Lastly, we evaluated the specificity of our sensor towards the SARS-CoV-2 spike S1 protein by comparing its real-time response against that of Middle East respiratory syndrome coronavirus (MERS-CoV) spike protein due to their genome similarities^66^. As can be seen in **Fig. 6c**, the tri-channel sensor can differentiate between the two proteins under physiological relevant conditions. For the MERS-CoV protein, the sensor shows no response with the signal remaining largely unaltered with increasing analyte concentration from 1 fM to 100 pM. On the contrary, exposure of the device to SARS CoV-2 spike S1 protein leads to a strong and systematic signal increase with increasing SARS-CoV-2 spike S1 protein concentration. The lowest concentration at which these differences are detectable can be deducted from **Fig. 6c** yielding a value of approximately 1 fM^56^. The LoD^67^ was estimated by applying the International Union of Pure and Applied Chemistry (IUPAC) protocol^68^ to the calibration plot for SARS-CoV-2 spike S1 protein in **Supplementary Figure 19c** yielding a value of 865 aM.

## Conclusions

We have developed a simple-to-manufacture, millimetre-scale, all-solid-state metal oxide transistor sensor that can detect the presence of biomolecules down to attomolar concentrations in real time while being operated under physiologically relevant environments. The unique device architecture combines high sensitivity and a large dynamic range in an all-solid-state sensing platform capable for analyte sensing in the liquid-phase. The versatile surface chemistry of the metal oxides employed allows for the incorporation of different receptor units (e.g. antibodies, enzymatic recognition elements, aptamers), which is anticipated to enable the detection of a broader range of biomolecules with extraordinary sensitivity and specificity. Furthermore, the ability to distinguish between negatively and positively charged biomolecules as well as between the SARS-CoV-2 and MERS-CoV spike proteins, showcases the universality of the sensor platform, which could be exploited for addressing the most urgent sensing applications.

## METHOD

### Preparation of metal-oxide precursors

ZnO and In_2_O_3_ precursor solutions were prepared by dissolving zinc oxide (99.99%; Sigma-Aldrich) in ammonium hydroxide (50% v/v; Alfa Aesar) at a concentration of 10 mg ml^-1^ and anhydrous indium nitrate (99.99%; Indium Corporation) in 2-methoxyethanol (99.8%; Sigma-Aldrich) at a concentration of 20 mg ml^-1^, respectively. As-prepared solutions were then stirred rigorously at room temperature for 24 h before use. This process yielded clear transparent oxide precursor solutions.

### Fabrication of low-dimensional oxide transistors

Heavily doped silicon (Si^++^) wafers with a thermally grown SiO_2_ top-layer (100 nm) were used as the common gate electrode and the gate dielectric, respectively. Prior to the semiconductor deposition, the substrates were sonicated in a solvent bath each lasting for ≈10 min in the following sequence: 1) deionised (DI) water with a Decon 90 detergent (5 vol%); 2) DI water; 3) acetone; 4) isopropanol. The solvent residue was dried with dry nitrogen over the substrate surface. As the last cleaning step, the substrates were exposed to ultraviolet (UV) ozone treatment for 10 min. The In_2_O_3_ ultra-thin film was deposited by carrying out spin-casting of the as-prepared precursor solution onto the Si substrates at 6000 rpm for 30 s in ambient air, followed by a post-deposition thermal-annealing process for 60 min at 200 °C in ambient air. The top ZnO layer was deposited with the same procedure as that for the In_2_O_3_ layer. Fabrication of the transistors (channel width/length = 1000/100 μm μ/m) was completed with thermal evaporation of 40-nm thick Al top source and drain (S–D) electrodes through a shadow mask in high vacuum (≈10^−6^ mbar).

### Transistor characterisation

Electrical characterisation of transistors was carried out using three micro-positioners (EB-700, EVERBEING), a homemade probe station and an Agilent B2902A source/measure unit in a nitrogen-filled glove box.

### Self-assembled layer preparation and surface modification

To prepare the modified device for DNA sensing, firstly 1-pyrenebutyric acid (PBA, 97%; Sigma-Aldrich) solution (1 mg ml^- 1^ in anhydrous tetrahydrofuran (THF)) was applied on the surface of the transistor for 30 min and thoroughly rinsed with THF and dried under nitrogen atmosphere. Butyric acid (BA, ≥ 99%; Sigma-Aldrich) solution (1 mg ml^-1^ in anhydrous THF) was then applied to the PBA modified surface for 30 min and thoroughly rinsed with THF and dried under nitrogen atmosphere. To prepare the modified device for avidin sensing, biotin (99%; Sigma-Aldrich) solution (0.8 mg ml^-1^ in anhydrous ethanol) was first applied on the surface of the transistor for 30 min and thoroughly rinsed with ethanol and dried under nitrogen atmosphere. BA was then applied to fully passivate the uncovered surface following the same procedures above as for DNA sensor device.

### Analyte preparation and sensing

Deoxyribonucleic acid from calf thymus (Type XV, Activated, lyophilised powder), avidin (lyophilised powder, ≥10 units/mg protein), A20, T20, (AT)20, were purchased from Sigma and used as received. All analytes were well dissolved in MilliQ water (18.2 MΩ·cm/25 °C) to reach the desired concentration according to the solution preparation instruction provided by the supplier. For the sensing process, the analyte solution was constantly applied onto the sensing area, and the electrical properties of the sensor devices were then recorded. For the real time sensing, the channel current was monitored during the continuous and consecutive application of analyte solution of different concentrations onto the same sensor device.

### Ultraviolet–visible spectroscopy measurements

The ultraviolet–visible (UV-Vis) transmission measurements were performed using a Shimadzu UV-2600 UV–Vis spectrophotometer. The samples were prepared on quartz substrates using the same deposition parameters described in the Methods section for oxide thin-film deposition and self-assembled monolayer formation.

### High-resolution transmission electron microscopy measurement

The samples for high-resolution transmission electron microscopy (HRTEM) analysis were prepared using the focused ion beam processing technique. A gold-plated layer with thickness of 5 nm was coated on sample via sputtering before the sample preparation to make its surface more conductive. The HRTEM images were acquired at 300 kV by a FEI Titan G2 80–300 microscope equipped with a high-brightness Schottky-field emission electron source and a high-resolution Gatan imaging filter Tridiem energy-filter.

### Atomic Force Microscopy measurement

Atomic force microscopy study was carried out in tapping mode using an Agilent 5500 atomic force microscope in ambient atmosphere. The approximate resonance frequency of the cantilever was 280 kHz and force constant was ≈60 N·m^-1^.

### Scanning Kelvin probe measurement

Scanning Kelvin probe investigations were carried out using a KP Technology system (model SKP5050/APS02) with a 1 mm tip. Scanning was achieved by taking an individual Kelvin probe (KP) measurement in one location and then moving the motorised stage to bring the sample in position for the next KP measurement. This was repeated until data was gathered in a grid pattern of 60×60 points, spanning an area of ca. 4 mm × 4 mm. For each point location the tracking feature built into the software made sure to keep the average tip-to-sample-distance constant. Additional drain bias in the range of 0 to 3 V was applied using a Keithley B2400 Source-Meter unit. The WF and E_F_ values were calculated using Silver as the reference material. All measurements were carried out in ambient air at room temperature and relative humidity of ca. 25 %.

### Real-time sensing data analysis of (AT)20, natural dsDNA from calf thymus and avidin

The sensors’ real-time recordings to synthetic dsDNA (AT)20 (**Fig. 4h**), natural dsDNA (**Fig. 5c**) and avidin (**Fig. 5f**) at different analyte concentrations were fitted according to the linear Langmuir adsorption isotherm equation^52^: C_analyte_/ΔI_D_ = C_analyte_/ΔI_sat_ + 1/ΔI_sat_K_eq_, where ΔI_sat_ is the change of the saturated channel current upon increasing the concentration of analyte, and K_eq_ is the binding constant of the analyte with its corresponding receptor.

### Solution preparation and device fabrication for spike S1 protein sensing experiments

The 3-aminopropyltriethoxylsilane (APTES) solution (99%), glutaraldehyde solution (70% in H_2_O), and phosphate buffered saline (PBS, pH 7.4, 10×) solution were purchased from Merck. SARS-CoV-2 spike S1 antibody (40150-R007), SARS-CoV-2 (2019-nCoV) Spike S1-His Recombinant Protein (40591-V08B1), MERS-CoV Spike/S1 Protein (S1 Subunit, aa 1-725, His Tag) (40069-V08H) were purchased from Sino Biological (China). All chemicals were used as received without further purification. Stock solutions of spike proteins were prepared using nuclease-free water and further diluted to different concentrations in 0.1× PBS where necessary. For the SARS-CoV-2 spike protein sensing, the tri-channel transistors were first treated with UV-irradiation for 10 min, APTES solution (2 wt%) in toluene was pipetted onto the oxide surface and left for 15 min, followed by rinsing with toluene and annealing at 120 °C for 1 h. A glutaraldehyde (GA) linker was added to the terminal amino (-NH_2_) groups of APTES using a solution of 0.8% GA in DI water for 10 min at room temperature, followed by rinsing with DI water and dried with a stream of N_2_ gas. Next, a spike antibody solution (200 μg mL^-1^) was applied onto the surface of the functionalised device and kept at room temperature for 5 h in order to immobilise the spike S1 antibodies via covalent bonding. To complete the immobilisation process, the devices were rinsed with 0.1× PBS to remove unbound antibodies. The presence of the antibodies on the surface of the sensing channel was verified via atomic force microscopy. The SARS-CoV-2 spike structure in **Fig. 6a** was adopted from the PDB ID: 6VYB^69^.

### Device modelling and simulations

The oxide transistor sensors were modelled and simulated using the semiconductor module in COMSOL Multiphysics. The cross-sectional model was constructed based on the actual device dimensions shown in **Supplementary Figure 3**. The oxide semiconductors were modelled based on material parameters taken from our previous reports^31,32,70^. The semiconducting In_2_O_3_/ZnO interface was modelled as a continuous quasi-Fermi-level heterojunction. The S-D electrodes were modelled as Ohmic contacts whilst the gate was modelled using the Thin Insulator Gate node, employing the same SiO_2_ dielectric condition as the actual device stack. The analyte was modelled as equivalent surface charges with a density of 10^10^ cm^-2^. All the other boundaries were modelled as insulations, indicating no normal flux such as current and electric displacement fields. Due to the large aspect ratio of the ultrathin oxide structures, the mapped mesh was generated for the entire transistor channel area with fine rectangular meshes.

## Data Availability

The data that support the plots within this paper and other findings of this study are available from the corresponding authors upon reasonable request.

## Acknowledgements

The authors would like to thank Prof. Arnab Pain, as well as Olga Douvropoulou and Raushan Nugmanova from the Biological and Environmental Science and Engineering (BESE) Division at KAUST (Saudi Arabia) for fruitful discussion and assistance with the materials related to the coronavirus spike protein sensing, and Dr Cheng Sheng Lin from Pitotech Co. Ltd (Taiwan) for useful suggestion and assistance in device modelling and simulation. P.P. and A.D.M. would like to acknowledge the postdoctoral funding for A.D.M. from Vidyasirimedhi Institute of Science and Technology (VISTEC). T.D.A. A.S., A.S., W.A., and H.F. acknowledge financial support from King Abdullah University of Science and Technology (KAUST) and KAUST Solar Centre.

## Author contributions

Y.-H.L., Y.H., M.H. and T.D.A. conceived the project. Y.-H.L. and Y.H. designed the experiments. Y.-H.L. designed and fabricated the tri-channel transistor sensors and thin-film samples, conducted the electrical characterisation, and analysed the device data. Y.H. prepared analytes, processed self-assembled monolayers and contributed to the molecular intercalation and kinetics analyses. W.S.A. and Abhinav S. performed additional sensing experiments and contributed to the analysis of the sensing data. Y.-H.L. and Y.H. performed the UV-Vis measurement. C.-H.L. performed device modelling with Y.-H.L., T.-H.C. and X.-W.X. and analysed the simulation results with Y.-H.L. P.P. and A.D.M. performed analyses on device parameters. H.F. carried out the scanning Kelvin probe measurement, collected the electrostatic potential data, assisted the analysis of the device data, and provided suggestion on the sensing measurement. Akmaral S. conducted the TEM characterisation. Y.-H.L., Y.H., M.H. and T.D.A. deduced the sensing mechanism and electronic process. T.D.A. supervised the whole project. All the authors discussed the results and contributed to the writing of the paper.

## ADDITIONAL INFORMATION

Correspondence and request for materials should be addressed to;

