## Supplementary data for "An all-solid-state heterojunction oxide transistor for the rapid detection of biomolecules and SARS-CoV-2 spike S1 protein"

**Supplementary Note 1 | Experimental and design details of the tri-channel transistor.**

The channel material for making a tri-channel transistor sensor is based on a modified $In_2O_3$/ZnO heterostructure. As-cleaned Si/$SiO_2$ substrates were first deposited with an ultrathin-layer of $In_2O_3$, followed by the deposition of a ZnO layer (see **Supplementary Figure 1** step 1-5). The fabrication protocols for preparing $In_2O_3$/ZnO precursor solutions and their depositions are the same as detailed in the Methods. The source-drain (S-D) electrodes were then formed onto these oxide double layers through thermal evaporation of 30-nm Al (**Supplementary Figure 1** step 6). To complete this modified $In_2O_3$/ZnO heterostructure, we first diluted the ZnO precursor (10 mg/mL in ammonium hydroxide) with isopropanol (IPA) to a concentration of 6 mg/mL and followed a similar ZnO deposition protocol (except for thermal annealing at 120 °C for 10 min) to form a layer of ZnO on top of the whole device area (**Supplementary Figure 1** step 7-8). The half-completed device was functionalised with 1-pyrenebutyric acid (PBA) through a self-assembly process by immersing the device in a PBA-containing solution for 30 min, followed by a thorough washing process with tetrahydrofuran (THF) and a brief post-annealing step for ~5 min at 50~60 °C (**Supplementary Figure 1** step 9~10). The same self-assembly process was then carried out for the formation of the butyric acid (BA) passivation layer (**Supplementary Figure 1** step 11-12). The preparation for the PBA and BA solution are detailed in the Methods. Regarding the sensing demonstration of biotin-avidin, the biotin functionalisation was carried out using the same self-assembly procedure as that for the PBA and BA formation. The exact design of our tri-channel transistor sensor is depicted in **Supplementary Figure 2**. The device is composed of four individual electrode fingers that together form three different channel areas. The sensing part sits right at the centre of the device with a square footprint spanning 4 $mm^2$ (2 mm by 2 mm) whilst two conventional transistor channels are located on both sides of the sensing area with a channel width of 1,800 μm and a channel length of 100 μm.

**Supplementary Note 2 | Impact on transconductance upon analyte exposure**

The transconductance, $g_m$, of a transistor is given by

$$g_m = \left. \frac{\partial I_D}{\partial V_G} \right|_{V_D} \tag{SI-1}$$

which is the rate of change of $I_D$ with respect to $V_G$. The transconductance entails several characteristics of the device: charge transport (mobility), capacitive coupling (geometric gate capacitance), geometrical factor (channel width and length), and operational parameters (gate and drain voltages). The complex geometry of the tri-channel transistor sensor coupled with the $In_2O_3$/ZnO HJ with quasi two-dimensional (q2D) charge distribution in the channel render the accurate analysis of carrier mobility difficult. In the following, we examine if $g_m$ is suitable for comparing the sensing performance, in particular when all sensors were based on an identical structure and operated under the same parameters.

The plots of transfer characteristics of tri-channel transistor sensors when exposed to (AT)20, A20, and T20 analytes on linear scale are shown in **Supplementary Figures 10a-c** (i.e. **Fig. 4b-d** in the main text but plotted on a linear scale). Linear relationships between $I_D$ and $V_G$ especially at high $V_G$ can be observed in all three plots. Since the devices were operated at $V_D$ = 3V, we fit a linear function to each of the transfer curves over a $V_G$ range of 20 V to 30 V. Ensuring that $V_D < V_G - V_{ON}$, i.e. the channel is in the linear regime, the slope of the fit is therefore the average linear-regime transconductance. Their normalised values are plotted in **Supplementary Figure 10d** to compare the changes in the transport properties as a function of analyte concentrations. As can be observed, the transconductance of the device increases with increasing (AT)20 and A20 concentrations. On the other hand, the transconductance decreases with that of T20, which may be due to Coulomb scattering[1]. However, the difference in $g_m$ for sensing A20 and T20 is highly doubtful to signify the sensitivity of our device as there are no known specific surface interactions or intercalations that would occur between pyrene and ssDNA. We note that although $\Delta I_D/g_m$ has been reported for the estimation of molecules adsorbed[2], the fact is that $g_m$ highly depends on what the 'regime' a transistor is operated on (e.g. deep-subthreshold, subthreshold, super-subthreshold, etc.) as well as how $g_m$ is extracted. Besides, $g_m$ is a parameter that requires calculation and cannot be output during measurement, hence much less valuable for direct generation of sensing data. Compared to the absolute current level, i.e. $I_D$, $g_m$ is a device parameter far less suitable for the interpretation of the sensing performance of a transistor sensor, and the bottom line is that a transistor needs to possess a adequate $g_m$ level to generate a perceivable change for signal distinguishability.

**Supplementary Note 3 | Responsivity and sensitivity for biochemical sensing**

We introduce two figures of merit: biochemical analyte responsivity ($R_{analyte}$) and analyte sensitivity ($S_{analyte}$)[3]. The $R_{analyte}$ is defined as the change in the channel current upon exposure to a specific analyte concentration [$I_D$(conc.) in relation to its initial baseline $I_D$ level, i.e. $I_D$(init.)], and is given as

$$R_{analyte} = \frac{I_D(\text{conc.}) - I_D(\text{init.})}{M(\text{conc.})} \quad \text{(SI-2)}$$

where $I_D$(init.) and $I_D$(conc.) are the drain current at a specific gate voltage before and after exposure to the analyte at each concentration, while M(conc.) is the analyte molar concentration. The $R_{analyte}$ (in amperes per molar or $A\cdot M^{-1}$) represents the output/input ratio defined as the gain of the sensor and provides information on the extent by which the response of the biosensor changes with varying analyte concentrations. **Supplementary Figure 11a** shows the evolution of $R_{analyte}$ with $V_G$ for A20, T20 and (AT)20 at different concentrations. For all analytes, the $R_{analyte}$ increases with increasing $V_G$, due to the transistor's intrinsic current amplification characteristics. The highest and lowest $R_{analyte}$ values are measured for the sensors exposed to (AT)20 and T20, respectively. Of particular interest is the extraordinary value of $R_{analyte}$ ($>10^{10}$ $A\cdot M^{-1}$) measured for the lowest concentrations of (AT)20, indicating that even at 100 aM the sensor is capable of inducing a change in current on the order of μA which can be easily detected. In some instances the value of $R_{analyte}$ decreases with increasing $V_G$ due to the sensing drain current [$I_D$(conc.)] approaching the initial channel current level [$I_D$(init.)]. This effect could be due to changes in the electric field distribution and electrostatic landscape upon changing the gate bias that may affect the surface interactions. Nevertheless, the sensor can be operated at different $V_G$ where the responsivity remains finite, demonstrating the operational versatility of employing a transistor as a sensing platform.

Although $R_{analyte}$ indicates gain, it does not account for the practicalities of operation. A sensor measures the change in signal above a baseline. The larger the background signal, the less accurate the signal measurement becomes. The analyte sensitivity $S_{analyte}$ provides a figure of merit that accounts for this by considering the change in current normalised to the background signal (i.e. initial current):

$$S_{analyte} = \frac{I_D(\text{conc.}) - I_D(\text{init.})}{I_D(\text{init.})} \qquad \text{(SI-3)}$$

In other words, the $S_{analyte}$ provides a measure for the sensor's reaction to the analyte by comparing the increase in the $I_D$ to the baseline current. **Supplementary Figure 11b** presents the $S_{analyte}$ versus applied $V_G$ and highlights the disparity in the sensor's response to A20, T20 and (AT)20. The dsDNA induces the largest response with a maximum value of 24,000. This extraordinary level of sensitivity is roughly five orders of magnitude larger than the values achieved in those state-of-the-art transistor sensors for comparable analyte concentrations[4,5]. Irrespective of analyte, the $S_{analyte}$ versus $V_G$ plots appear almost identical exhibiting a peak at around $V_G = V_{ON}$(init.).

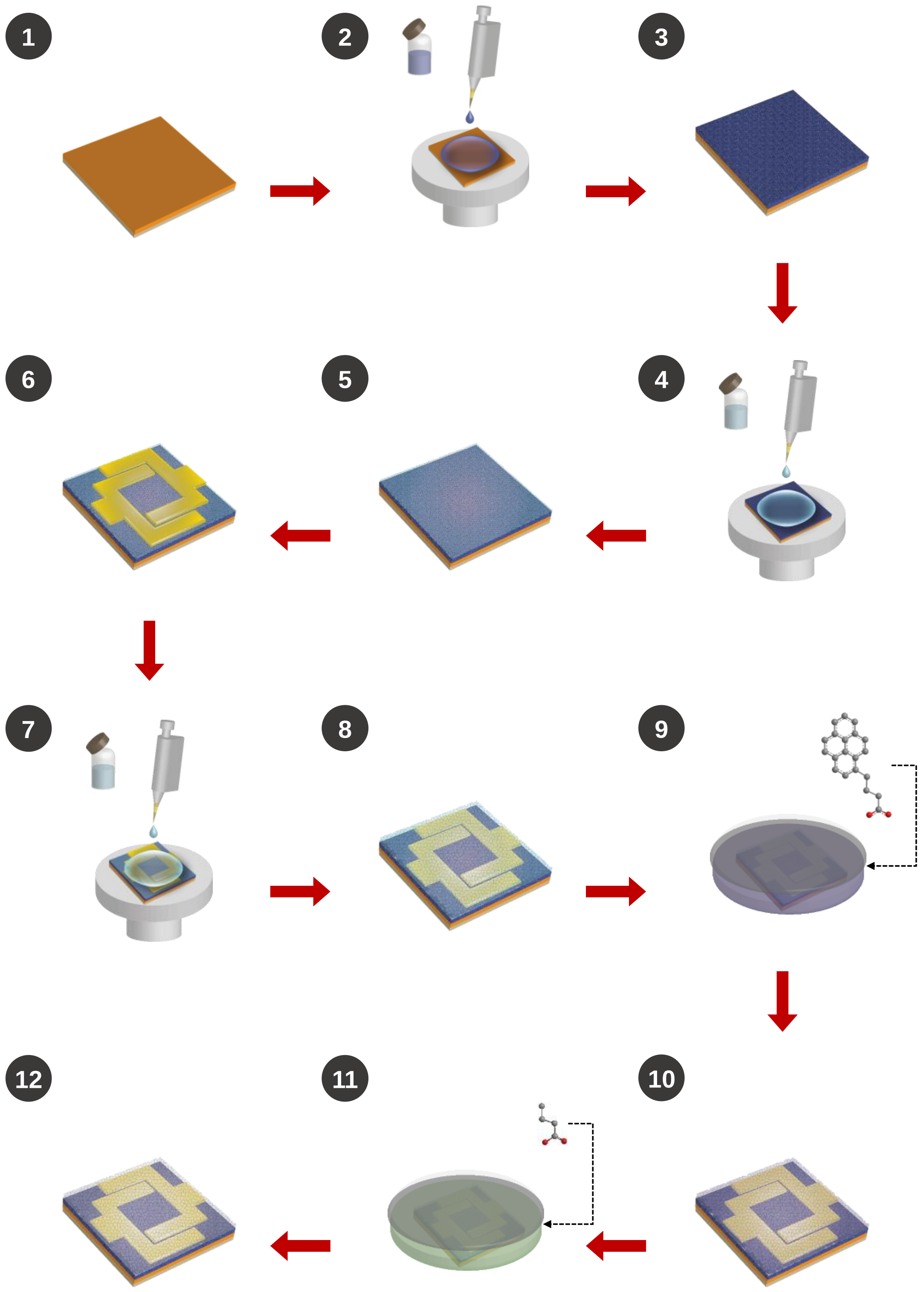


**Supplementary Figure 1 | Fabrication process of tri-channel transistor sensor.** Schematic of the fabrication process: 1-3) deposition and formation of the bottom $In_2O_3$ layer onto a $Si/SiO_2$ substrate; 4-5) deposition and formation of the $1^{st}$ ZnO layer; 6) Al top electrodes deposition; 7-8) deposition and formation of the $2^{nd}$ ZnO layer; 9) PBA SAM formation; 10-11) BA SAM formation; 12) complete device.

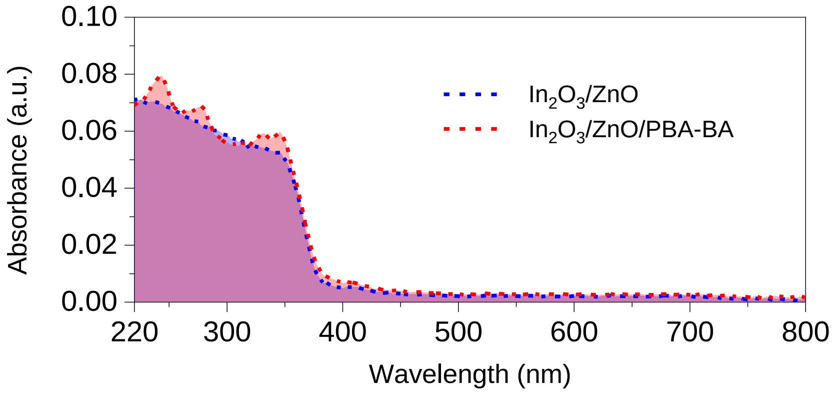


**Supplementary Figure 2 | Ultraviolet–visible absorption spectrum.** UV-Vis absorption measurements taken for the intrinsic $In_2O_3$/ZnO hetero film stacks before and after applying the PBA and BA self-assembly processes.

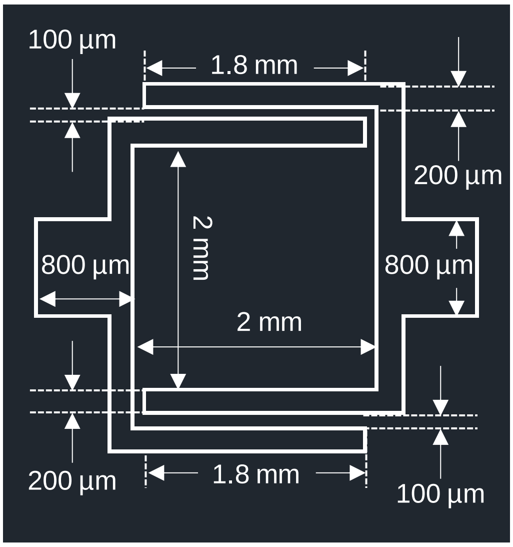


**Supplementary Figure 3 | Design of tri-channel transistor sensor.** A tri-channel transistor sensor composed of three individual channels: 1) one sensing channel with a square footprint spanning 2 mm by 2 mm in the centre; 2) two identical conventional transistor channels on the sides with a channel width/length of 1,800/100 µm/µm.

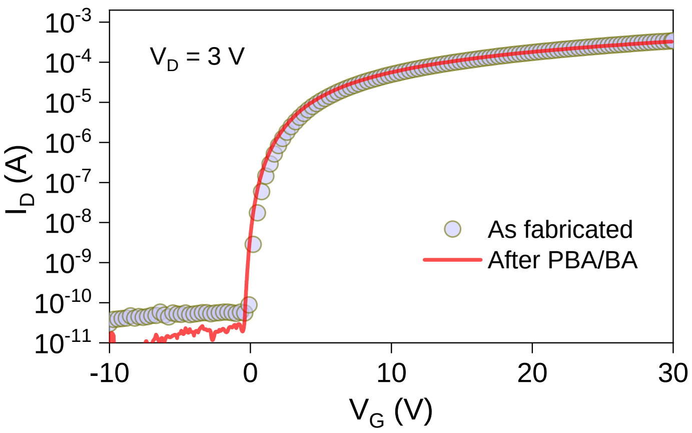


**Supplementary Figure 4 | Transfer characteristics before/after SAM functionalisation.** Transfer current-voltage (I-V) characteristics of a tri-channel transistor sensor measured as fabricated and after the PBA/BA self-assembly processes.

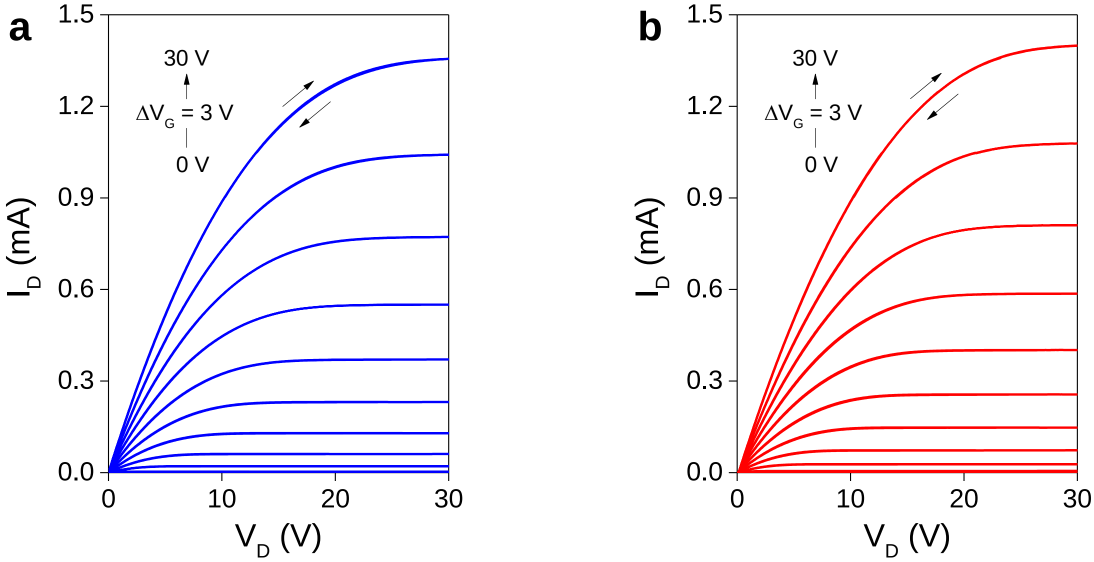


**Supplementary Figure 5 | Output characteristics before/after SAM functionalisation. a,b,** Output current-voltage (I-V) characteristics of a tri-channel transistor sensor measured (**a**) as fabricated and (**b**) after the PBA/BA self-assembly processes.

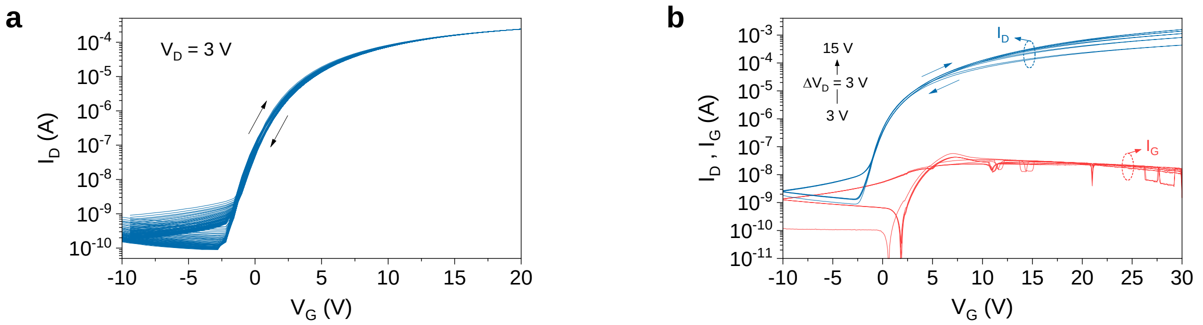


**Supplementary Figure 6 | Repeated transfer characteristics and gate leakage current. a**, Representative 90 forward-backward dual sweeps of transfer I-V characteristics measured from a tri-channel transistor. **b**, A typical set of transfer characterisation results for a tri-channel transistor sensor exhibits negligible contribution from gate leakage current to device electrical performance beyond $V_{ON}$.

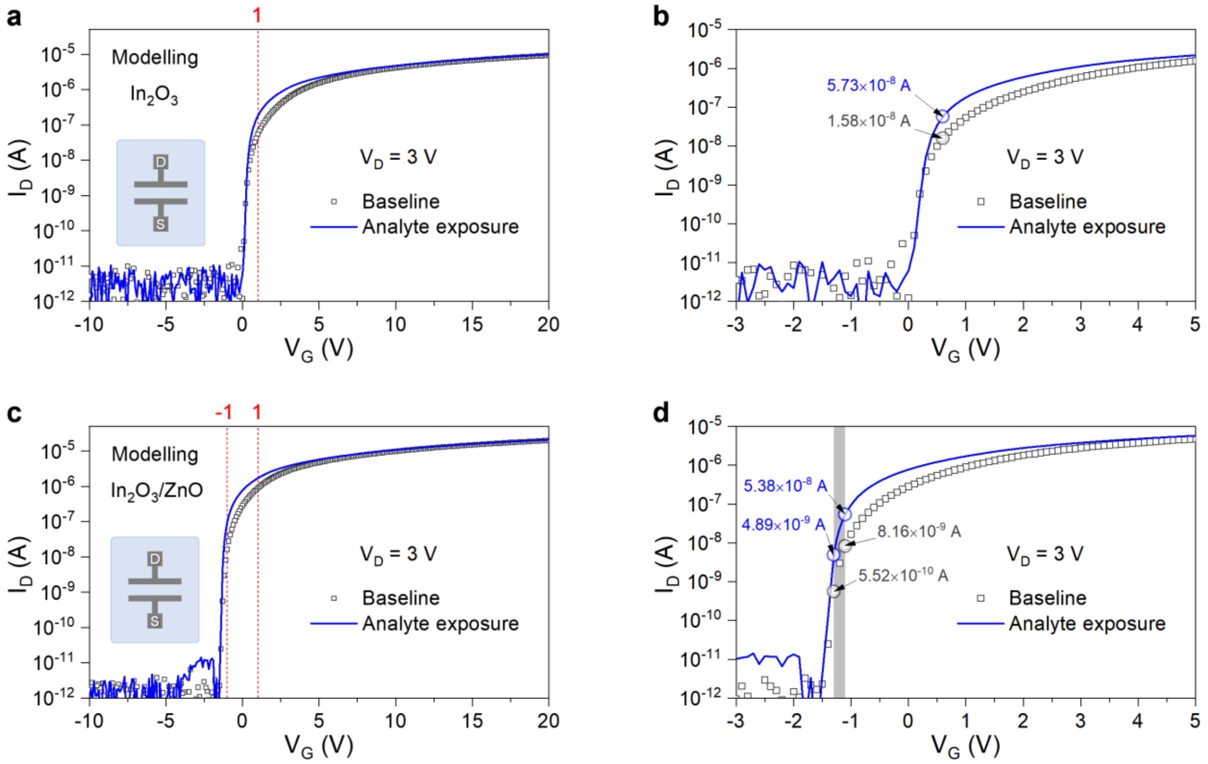


**Supplementary Figure 7 | Modelled current-voltage characteristics for oxide transistors based on the conventional channel architecture. a**,**b**,**c**,**d**, COMSOL modelled transfer current-voltage (I-V) characteristic obtained at $V_D$ = 3 V. Modelled transfer characteristics for transistors based on a single channel layer of $In_2O_3$ (**a**-**b**) and an $In_2O_3$/ZnO heterojunction (**c**-**d**). (**a**) and (**c**) show the full ranges of simulated I-V characteristics whilst (**b**) and (**d**) show the I-V characteristics for $V_G$ in the range -3 to 5 V. When comparing the largest differences in the channel current ($\Delta I_D$) for baseline and exposure to a simulated analyte, the largest $\Delta I_D/I_0$ obtained for $In_2O_3$/ZnO and $In_2O_3$ transistors are 7.86 and 2.63, respectively.

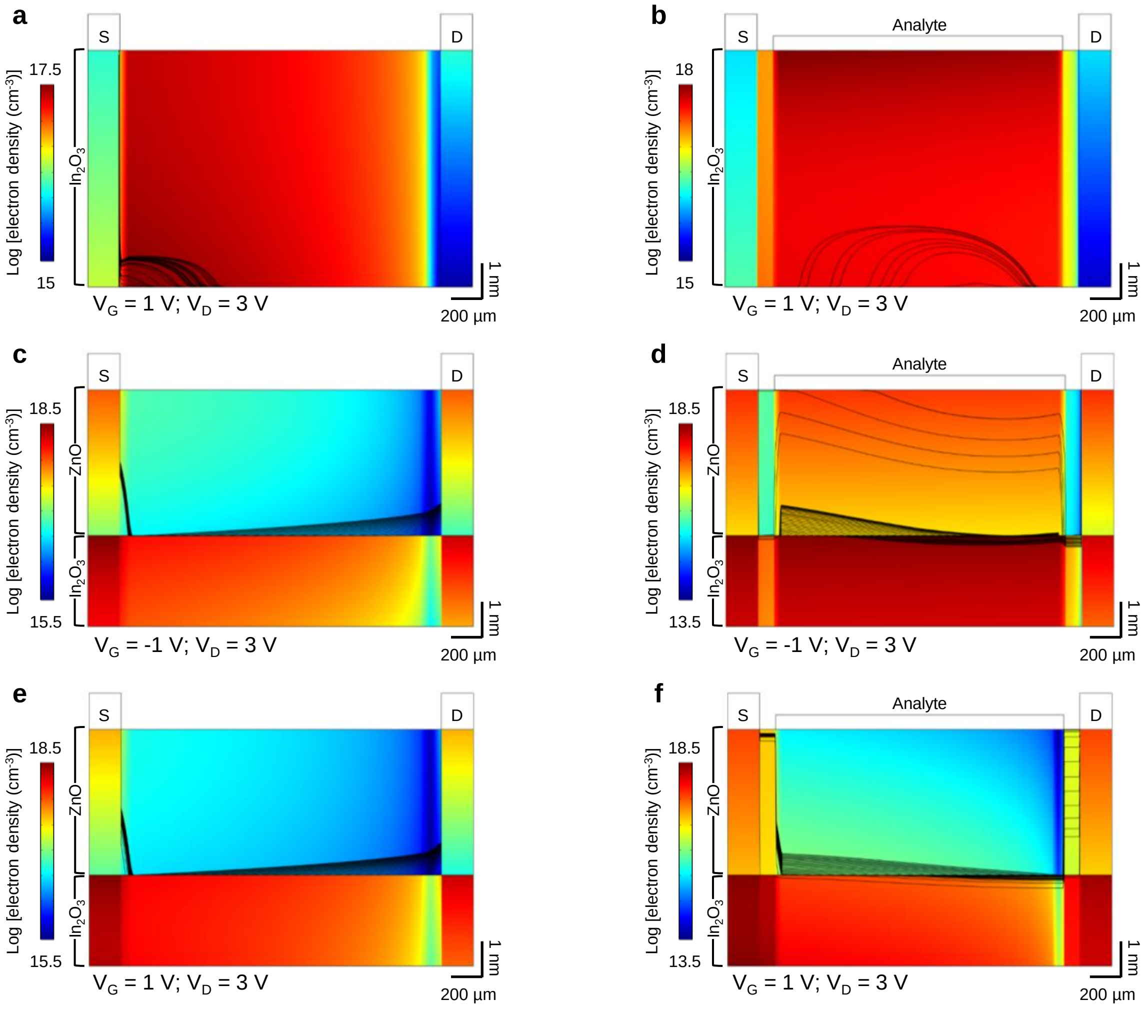


**Supplementary Figure 8 | Simulated electron density distributions. a**,**b**,**c**,**d**,**e**,**f**, COMSOL modelled electron density distributions and electron flow streamlines (all under $V_D$ = 3 V) for baseline: (**a**) $In_2O_3$ ($V_G$ = 1 V), (**c**) $In_2O_3$/ZnO ($V_G$ = -1 V) and (**e**) $In_2O_3$/ZnO ($V_G$ = 1 V); and under the exposure of simulated analytes (**b**) $In_2O_3$ ($V_G$ = 1 V), (**d**) $In_2O_3$/ZnO ($V_G$ = -1 V) and (**f**) $In_2O_3$/ZnO ($V_G$ = 1 V). The electrodes and analytes are shown to indicate their positions with respect to the devices.

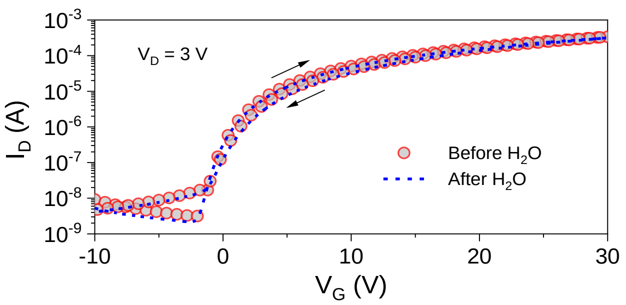


**Supplementary Figure 9 | Transfer characteristics of sensor under the presence of $H_2O$.** Transfer I-V characterisation for a tri-channel transistor sensor carried out with $H_2O$ present on the device sensing area. There is virtually no change in the key performance parameters: as for the initial state, $V_{ON}$ = -2.0 V, $I_{ON}$ = 0.32 mA, S.S. = 0.72 V/dec; and under the presence of $H_2O$, $V_{ON}$ = -2.1 V, $I_{ON}$ = 0.32 mA, S.S. = 0.73 V/dec.

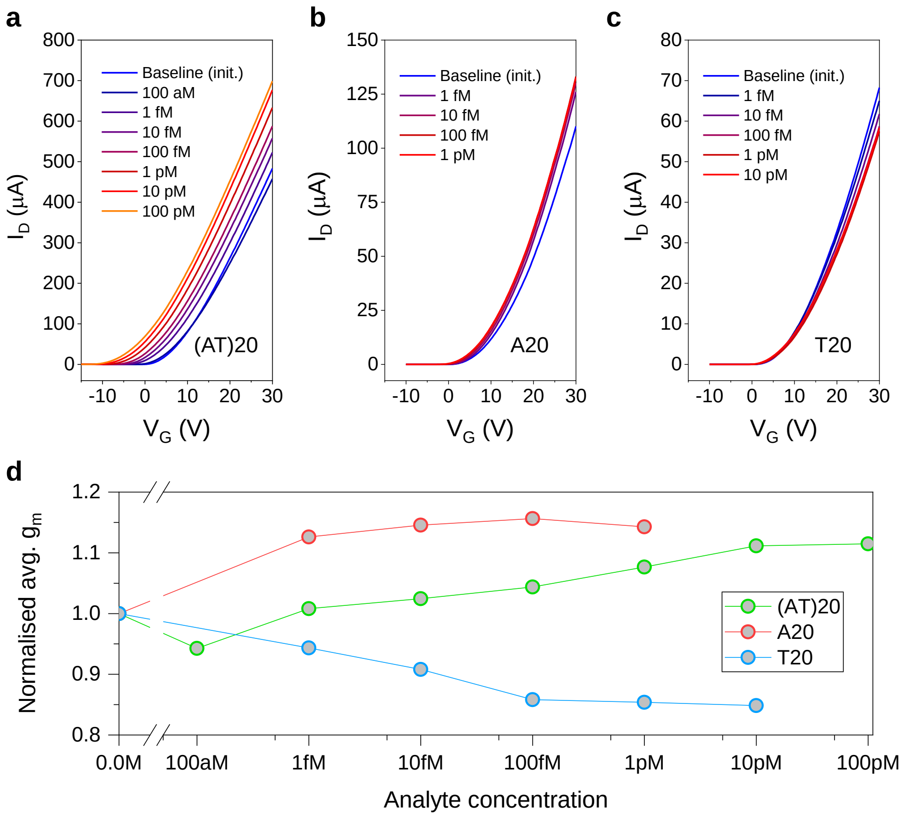


**Supplementary Figure 10 | Linear transfer characteristics and transconductance. a,b,c,** Plots of $I_D$ vs. $V_G$ in linear scale of tri-channel transistor sensors at various concentrations of: (a) (AT)20, (b) A20, and (c) T20. The linear relationship particularly at high $V_G$ shows that the devices operate in the linear regime. **d,** Average linear transconductance of the plots in (a)-(c) calculated from the slope of the linear portion in the range $V_G$ = 20-30 V. The values are normalised to the initial baseline values (no analyte).

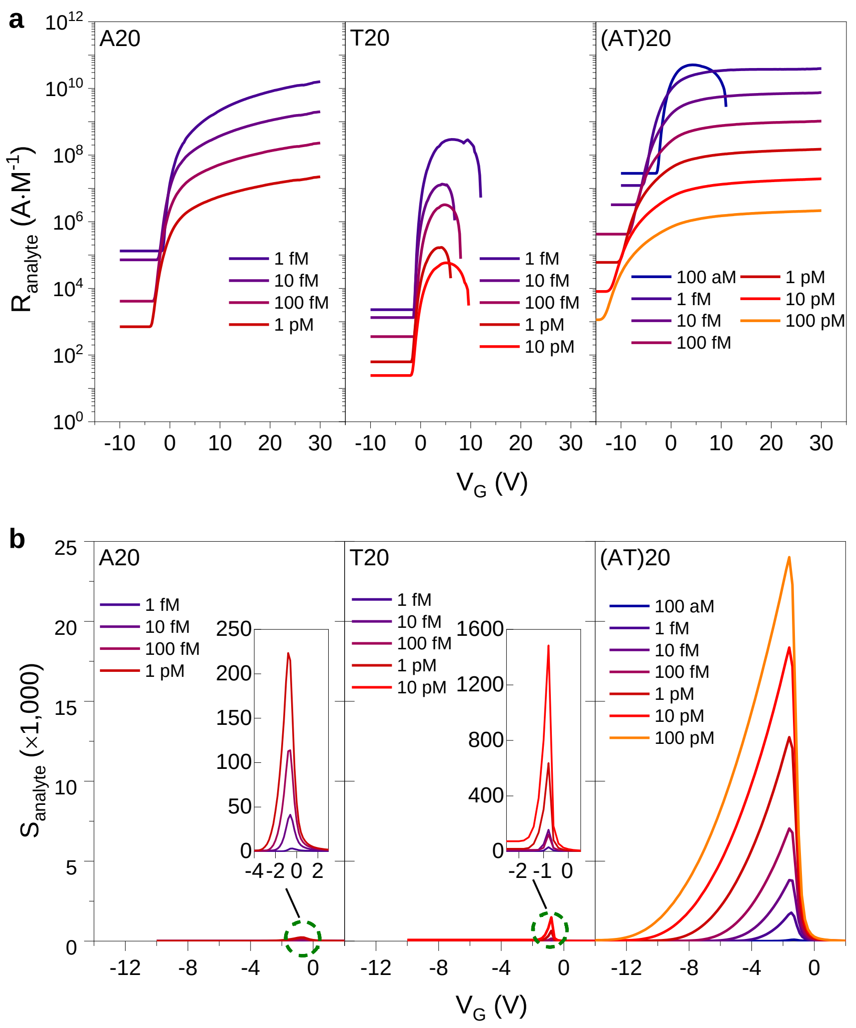


**Supplementary Figure 11 | Evaluation of the tri-channel transistor sensor performance. a**, Plots of analyte responsivity $R_{analyte}$ versus applied gate voltage for the different analyte DNAs studied at various concentrations. $R_{analyte}$ represents the $V_G$-dependent output/input signal gain of the biochemical-sensing. **b**, Plots of the analyte sensitivity $S_{analyte}$ as a function of gate voltage for the different DNA analytes at different concentrations. $S_{analyte}$ represents the signal-to-noise ratio of the biochemical-sensing transistor.

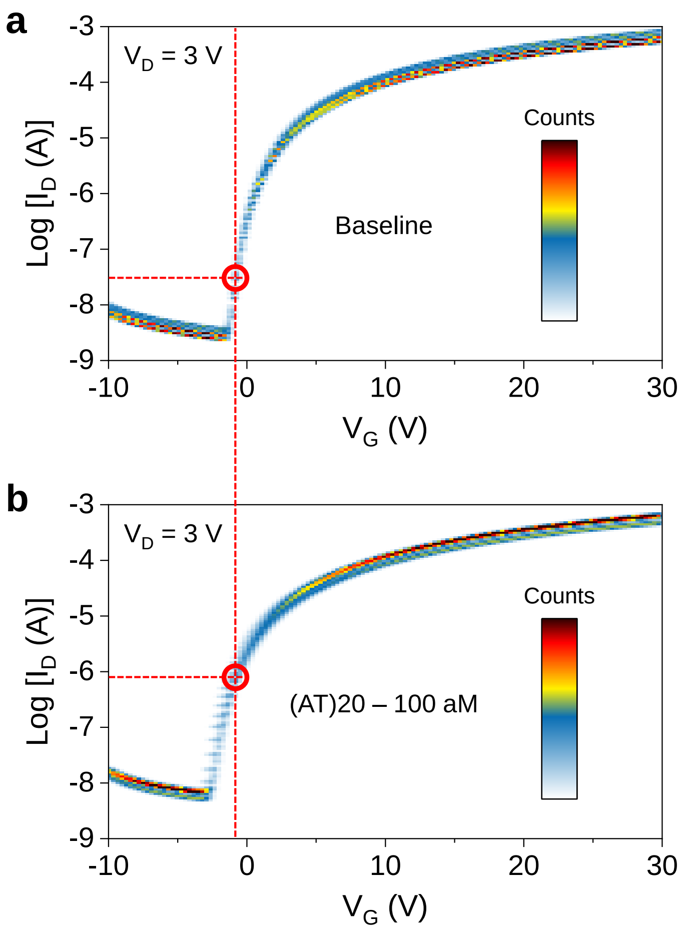


**Supplementary Figure 12 | Transfer characteristics without/with the presence of (AT)20.** Density plots of transfer I-V characteristics measured from 8 individual tri-channel transistors for (**a**) baseline and under (**b**) 100 aM of (AT)20. Red dashed lines indicate the $I_D$ levels at $V_G$ = -1 V.

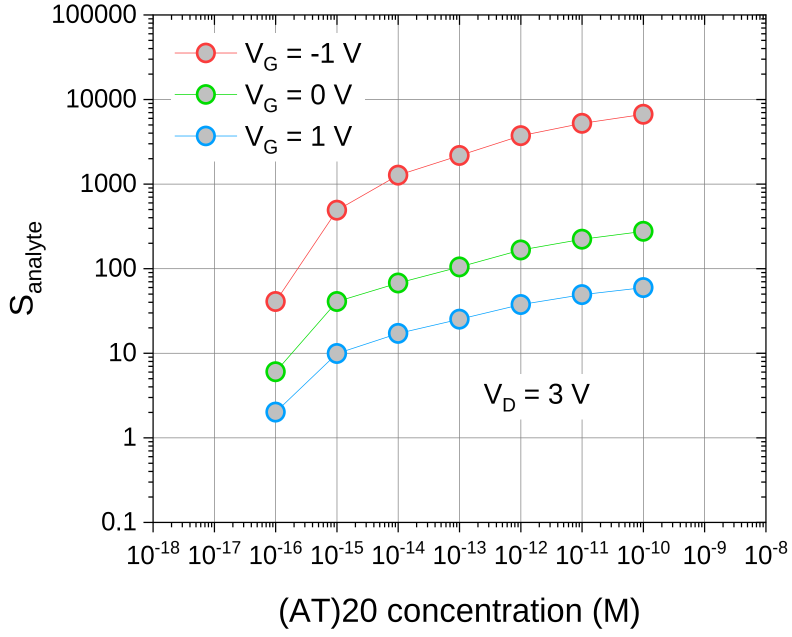


**Supplementary Figure 13 | Analyte sensitivity for (AT)20 in different operating regimes.** Plots of $S_{analyte}$ vs (AT)20 at a range of concentrations from 100 aM to 100 pM when operating a tri-channel transistor sensor at $V_D$ = 3 V and $V_G$ = -1 V (red), 0 V (green) and 1 V (blue). Higher $S_{analyte}$ achieved from biasing at $V_G$ = -1 V is a result of the low initial/baseline channel current $I_D$. When operating at $V_G$ = 1 V, the increase in the initial $I_D$ level reduces the distinguishable margin in $I_D$ for the lowest analyte concentration (i.e., 100 aM). However, one should also note that even though the lowest $S_{analyte}$ value of ~2 measured from the biasing condition of $V_D$ = 3 V and $V_G$ = 1 V, our solid-state tri-channel transistor sensor still outperforms current state-of-the-art transistor sensors reported in the literature using electrochemical approaches at similar analyte concentrations[4,6].

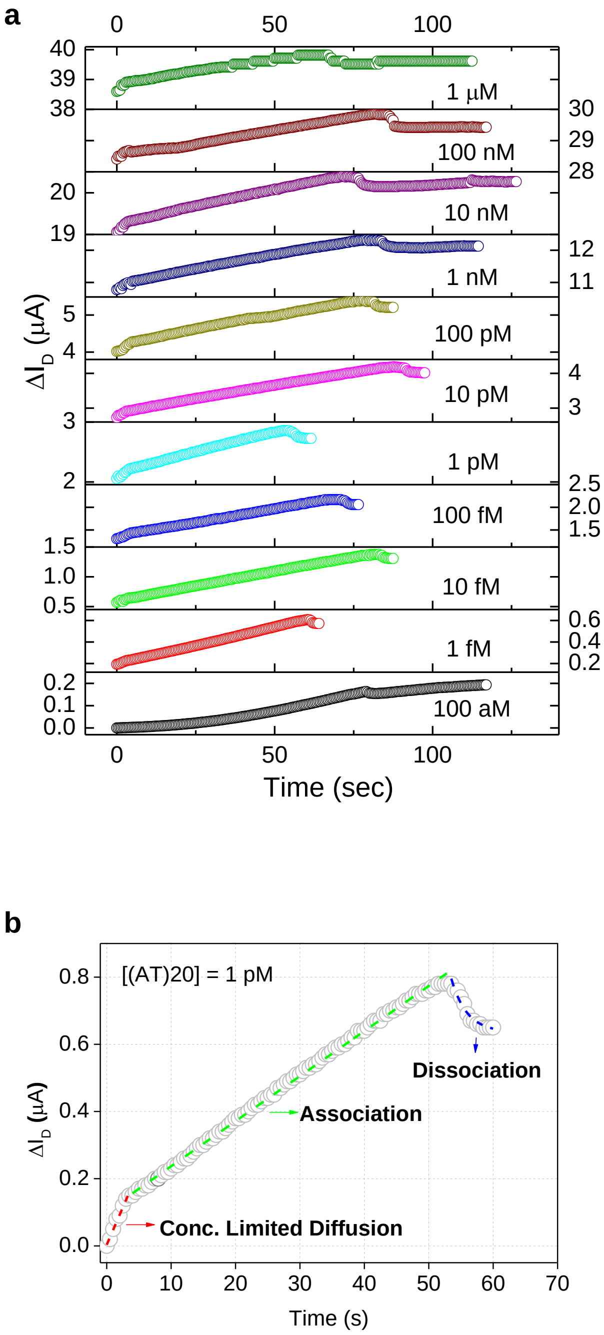


**Supplementary Figure 14 | Time course of the sensing process of (AT)20. a**, Monotonic increase in channel current at different (AT)20 concentrations. **b**, An example time course at [(AT)20] = 1 pM. The entire sensing process can be divided into three steps: a short time concentration-limited diffusion (the diffusion is faster when a higher (AT)20 concentration is applied); the association of (AT)20 with pyrene moieties on the surface of the sensor device via intercalation (the main sensing step, a zero-order reaction); and the dissociation of the formed complex until a thermal dynamic equilibrium is reached. The concentration-limited diffusion was fitted with a linear model: $y = 0.000344 + 0.00461x$ ($R^2 = 0.985$). The zero-order association step was fitted based on a linear model of $y = 0.0107 + 0.0014x$ ($R^2 = 0.998$). The dissociation step was fitted according to an exponential decay: $y = 0.0653 + 0.0163 \times \exp[-(x-53.5) / 2.65]$ ($R^2 = 1.00$).

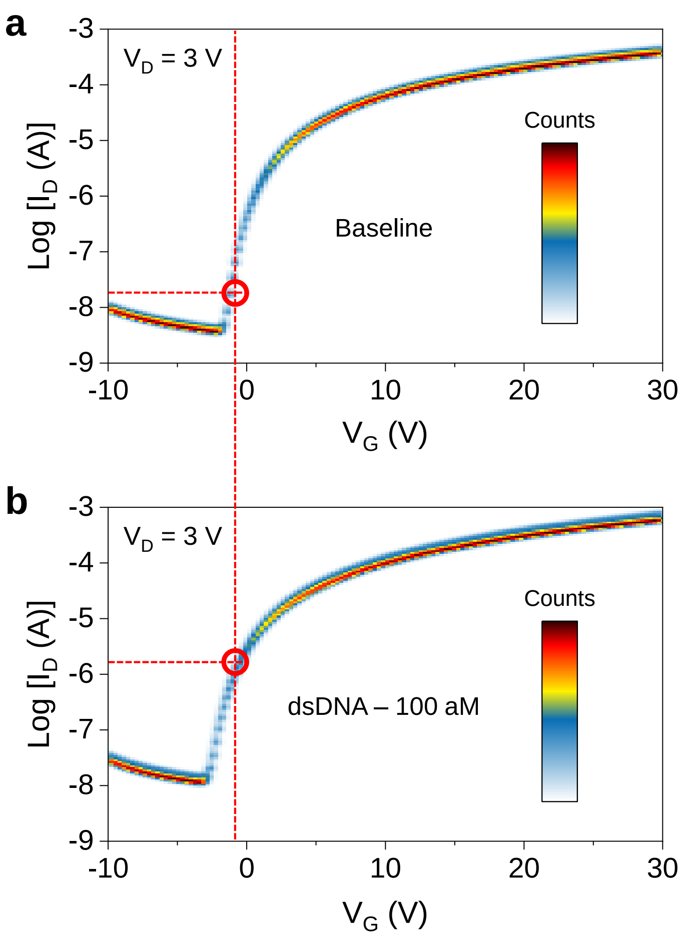


**Supplementary Figure 15 | Transfer characteristics without/with natural dsDNA.** Density plots of transfer I-V characteristics measured from 8 individual tri-channel transistors for (**a**) baseline and under (**b**) 100 aM of natural dsDNA. Red dashed lines indicate the $I_D$ levels at $V_G$ = -1 V.

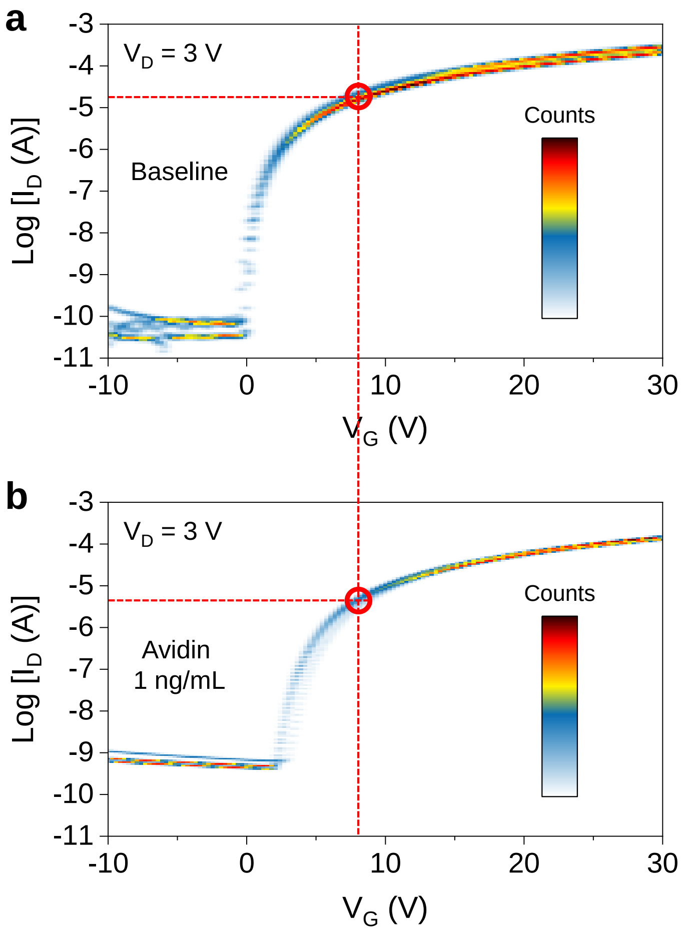


**Supplementary Figure 16 | Transfer characteristics without/with the presence of avidin.** Density plots of transfer I-V characteristics measured from 8 individual tri-channel transistors for (**a**) baseline and under (**b**) 1 ng/mL of avidin. Red dashed lines indicate the $I_D$ levels at $V_G$ = 8 V.

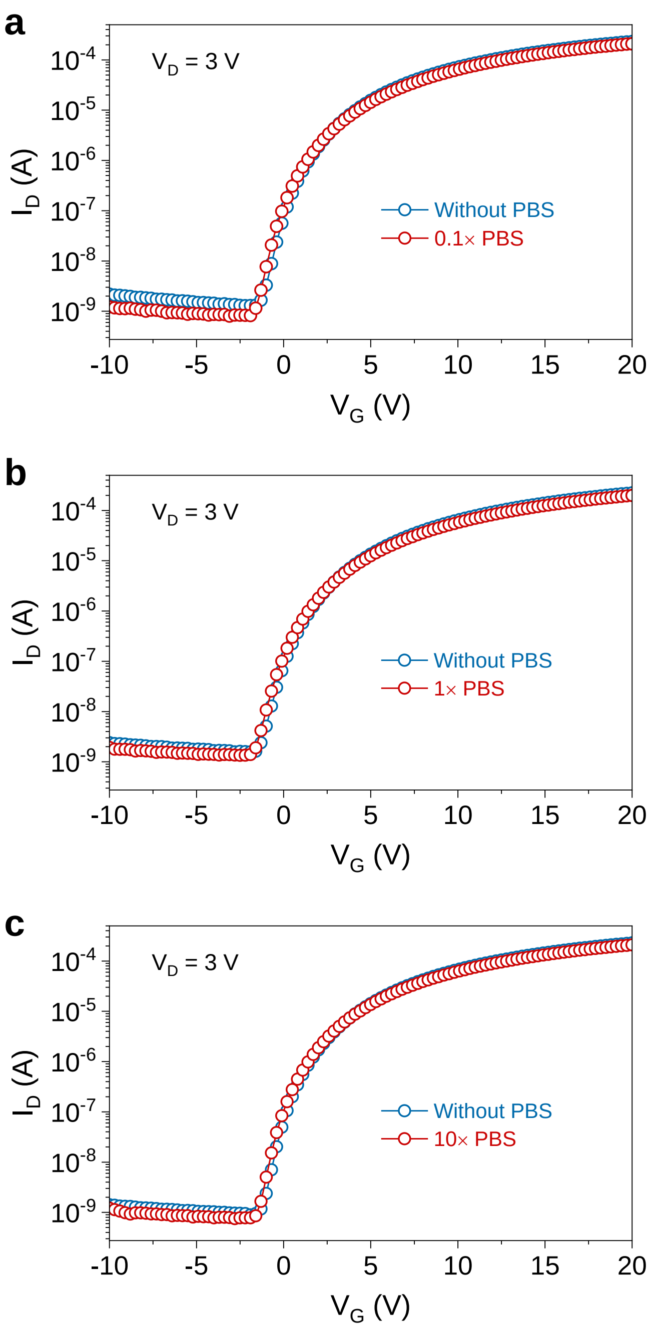


**Supplementary Figure 17 | Transfer characteristics without/with the presence of PBS.** Transfer I-V characteristics of tri-channel transistor sensors measured without and with the presence of (**a**) 0.1× PBS; (**b**) 1× PBS; (**c**) 10× PBS.

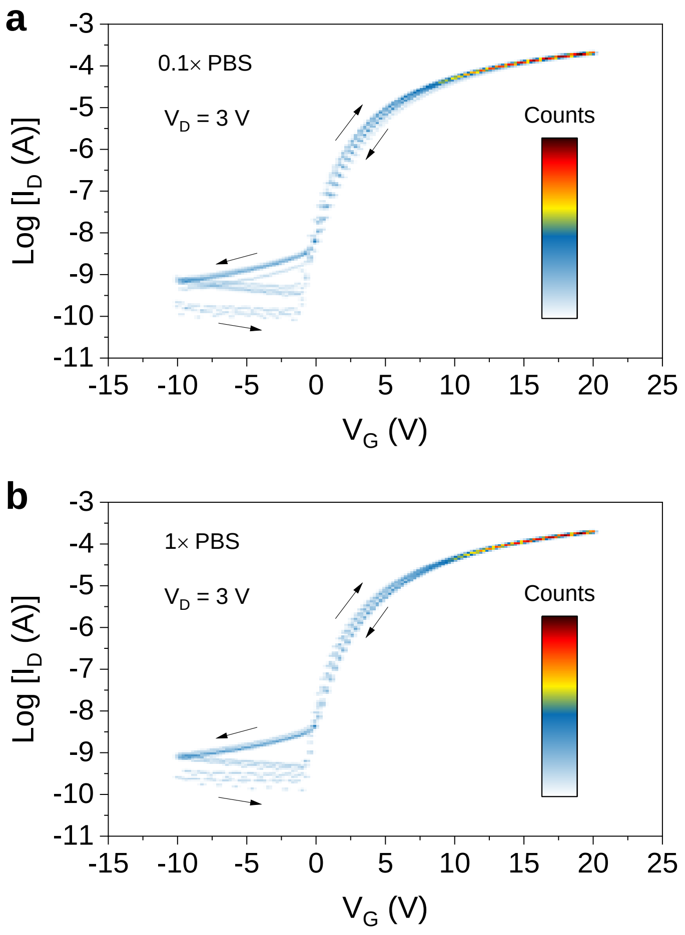


**Supplementary Figure 18 | Transfer characteristics under the presence of PBS.** Density plots of 5 sets of transfer I-V characteristics from tri-channel transistors under (**a**) 0.1× and (**b**) 1× PBS.

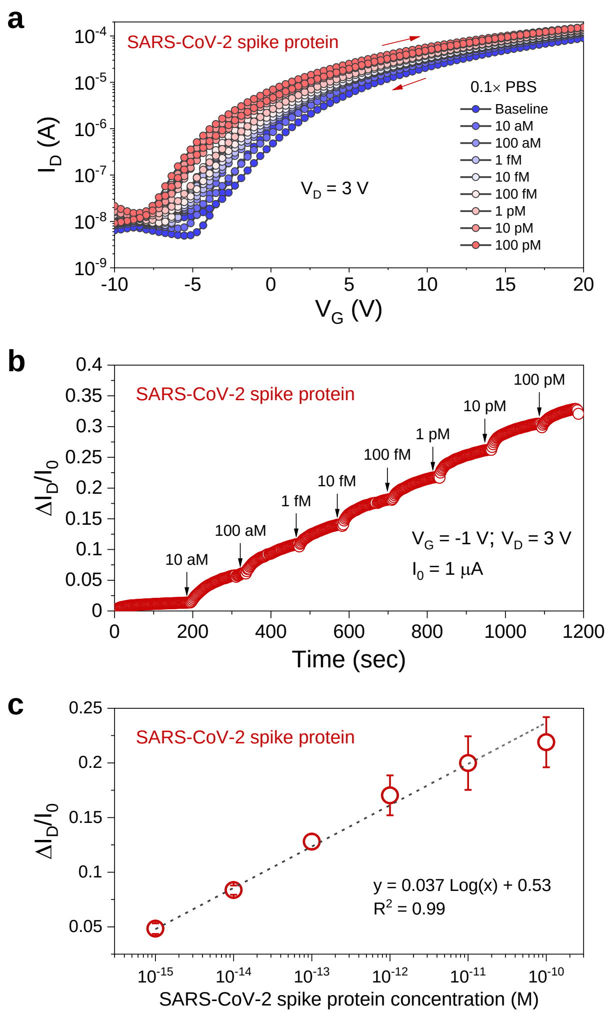


**Supplementary Figure 19 | Detection of SARS-CoV-2 spike protein. a**, Dual sweep transfer I-V characteristics for a tri-channel transistor sensor subjected to a series of SARS-CoV-2 spike protein concentrations in 0.1× PBS (baseline). **b**, Representative real-time detection of SARS-CoV-2 spike protein in 0.1× PBS. **c**, Corresponding calibration plot from (**b**). The error bars denote standard deviations from three real-time measurement sets. The limit of detection of 865 aM was estimated from 3 $S_b$/m, where $S_b$ is the standard deviation of the measurements for blank PBS-only solution, and m is the slope of the calibration curve[7].
